# Anti-tumor effects of an Id antagonist with no acquired resistance

**DOI:** 10.1101/2020.01.06.894840

**Authors:** Paulina M. Wojnarowicz, Marta Garcia Escolano, Yun-Han Huang, Bina Desai, Yvette Chin, Riddhi Shah, Sijia Xu, Ouathek Ouerfelli, Rajesh Kumar Soni, John Philip, David C. Montrose, John H. Healey, Vinagolu K. Rajasekhar, William A. Garland, Larry Norton, Neal Rosen, Ronald C. Hendrickson, Xi Kathy Zhou, Antonio Iavarone, Joan Massague, Andrew J. Dannenberg, Anna Lasorella, Robert Benezra

## Abstract

Id proteins are helix-loop-helix (HLH) transcriptional regulators frequently overexpressed in cancer. Id proteins inhibit basic HLH transcription factors through protein-protein interactions, often inhibiting differentiation and sustaining proliferation. We recently identified a small-molecule, AGX51, which targets Id proteins for degradation and impairs ocular neovascularization in mouse models. Here we show that AGX51 treatment of cancer cell lines impaired cell growth and viability that results from a dramatic increase in ROS production upon Id degradation. In mouse models, AGX51 treatment suppressed breast cancer colonization in the lung, regressed the growth of paclitaxel-resistant breast tumors when combined with paclitaxel and reduced tumor burden in a model of sporadic colorectal neoplasia. Furthermore, in cells and mice, we failed to observe acquired resistance to AGX51 likely the result of the immutability of the binding pocket and efficient degradation of the Id proteins. Thus, AGX51 is a first-in-class compound that antagonizes Id proteins, shows strong anti-tumor effects and may be further developed for the management of multiple cancers.

## Introduction

The Id proteins, Id1, Id2, Id3, and Id4, are helix-loop-helix (HLH) transcriptional regulators. They function by binding to and sequestering basic HLH (bHLH) transcription factors (e.g. the E protein E47). Id proteins are expressed during development and inhibit bHLH transcription factors to block differentiation and maintain self-renewal. Id protein expression is largely silenced in adult tissues but can be re-activated in several disease processes, including cancer (Lasorella et al., 2014; Ling et al., 2014) (also reviewed in (Wang and Baker, 2015)). Initial data associating Id1 and Id3 with cancer emerged from studies of xenografts and spontaneous tumors in genetically engineered mouse models, which typically showed decreased tumor growth and impaired angiogenesis in *Id1* and/or *Id3* deficient backgrounds (Lasorella et al., 2014; Lyden et al., 1999; Ruzinova et al., 2003). ID1 and ID3 are highly expressed in virtually all human cancers, in the vasculature and/or the tumor cells, including solid tumors of the breast (Perk et al., 2006; Schoppmann et al., 2003; Wazir et al., 2013), pancreas (Lee et al., 2004), bladder (Perk et al., 2006), cervix/uterus (Alcantara Llaguno et al., 2009; Maw et al., 2008; Schindl et al., 2001), colon (Zhang et al., 2015; Zhao et al., 2008), endometrium (Takai et al., 2001; Takai et al., 2004), stomach (Han et al., 2004; Yang et al., 2011), nervous system (Anido et al., 2010; Lyden et al., 1999; Vandeputte et al., 2002), liver (Matsuda et al., 2005), ovary (Schindl et al., 2003), prostate (Coppe et al., 2004; Forootan et al., 2007; Sharma et al., 2012), kidney (Li et al., 2007), esophagus (Liu et al., 2010), lung (Castanon et al., 2013; Kim et al., 2013; Ponz-Sarvise et al., 2011), and thyroid (Ciarrocchi et al., 2011; Kebebew et al., 2004) as well in hematopoietic tumors such as AML (Tang et al., 2009). In virtually all of these cancer types the presence of Id1 and/or Id3 is associated with an aggressive phenotype and poor clinical outcome. However, mutations in the genes or their promoters are rarely found. One notable exception is Burkitt’s lymphoma where *ID3* is frequently mutated and appears to act as a tumor suppressor, perhaps due to the growth promoting properties of E proteins in these cells (Richter et al., 2012).

Id overexpression in cancer is largely due to the convergence of several pro-proliferative and pro-oncogenic signaling cascades on the Id promoters such as MAP kinase (Tournay and Benezra, 1996(Cook et al., 2016), Myc (Lasorella et al., 2000), BMPs (Ogata et al., 1993; Ying et al., 2003), Src (Gautschi et al., 2008), FLT3 (Tang et al., 2009), VEGF (Ciarrocchi et al., 2007), Tgf-β (Kang et al., 2003), Wnt (Rockman et al., 2001) and Notch signaling (Liu and Harland, 2003). High Id expression is associated with increased metastatic potential. For example, in breast cancer, high levels of ID1, ID3 and ID4 are present in the tumor cells of the highly metastatic, triple negative subtype (Gupta et al., 2007; Wen et al., 2012). In metastatic tumor cells, Id1 and Id3 facilitate the mesenchymal-to-epithelial conversion required for macro-metastatic disease progression, likely through antagonism of the bHLH protein Twist (Stankic et al., 2013).

We recently identified a small-molecule antagonist of Id proteins based on X-ray crystal structure coordinates and an *in silico* screen (Wojnarowicz et al., 2019). The top candidate from this screen, AGX51, caused a change in the secondary structure of purified Id1 and inhibited the ability of Id1 to associate with its target E proteins in cells. This in turn led to the destabilization of Id1 via ubiquitin-mediated proteolysis. All 4 members of the Id family were similarly destabilized by AGX51. Consistent with the low level expression of Id proteins in normal adult tissues, high doses of the drug were well tolerated in mice with no apparent toxicity (see below). Here we report the characterization of AGX51 activity in several cancer cell lines, patient derived xenografts (PDXs), pancreatic ductal adenocarcinoma (PDA) organoid cultures and a chemically-induced murine model of colorectal adenoma.. We demonstrate that Id inhibition by AGX51 impairs cell viability, increases ROS levels and inhibits tumor growth and cancer cell colonization of the lungs in mice, thus phenocopying the effects seen in genetic animal studies. Importantly, long-term exposure of multiple cancer cell lines to AGX51 failed to demonstrate evidence of acquired resistance likely due to the binding of the drug to a region of the proteins that is largely immutable.

## Results

### Effects of AGX51 on Id proteins

We recently reported that AGX51 treatment of HUVEC and HCT116 cells leads to disruption of the Id-E protein-protein interaction (PPI), resulting in ubiquitination of Id1 and its subsequent degradation (Wojnarowicz et al., 2019). To further characterize the effects of AGX51 in cancer cells, we treated the 4T1 murine mammary cancer cell line with increasing concentrations of AGX51 (0-80 μM) for 24 hours and observed a significant decrease in Id1 protein levels starting at 40 μM (Figure 1A). 4T1 cells treated with 40 μM AGX51 for 0-72 hours showed a decrease in Id1 levels starting at 4 hours, with near complete Id1 loss by 24hours (Figure 1B, Figure S1A). Id3 and Id4 are also lost albeit with delayed kinetics. Id2 is not expressed in 4T1 cells (data not shown) but we previously showed that ID2 levels decrease in response to AGX51 treatment in HCT116 cells (Wojnarowicz et al., 2019). Similar to 4T1 cells, HMLE *RAS Twist* cells (immortalized human mammary epithelial cells overexpressing Twist) also showed reduced levels of ID1, ID3 and ID4 when treated with 20 μM AGX51 over similar time courses (Figure S1B). A decrease in ID1 and ID3 protein levels was also observed in a patient derived xenograft (PDX) cell line (IBT) generated from a patient with triple negative breast cancer (TNBC) (Figure 1C; Figure S1C). The PDX cell line does not express ID2 and ID4 (data not shown). To determine the duration of Id1 and Id3 protein loss, 4T1 cells were treated with 40 μM AGX51 for 24 hours, then cultured in AGX51-free media over 48 hours. Id1 protein levels increased over time, reaching untreated levels by 24 hours (Figure 1D). Id3 protein levels showed slower recovery (Figure 1D).

**Figure 1.**
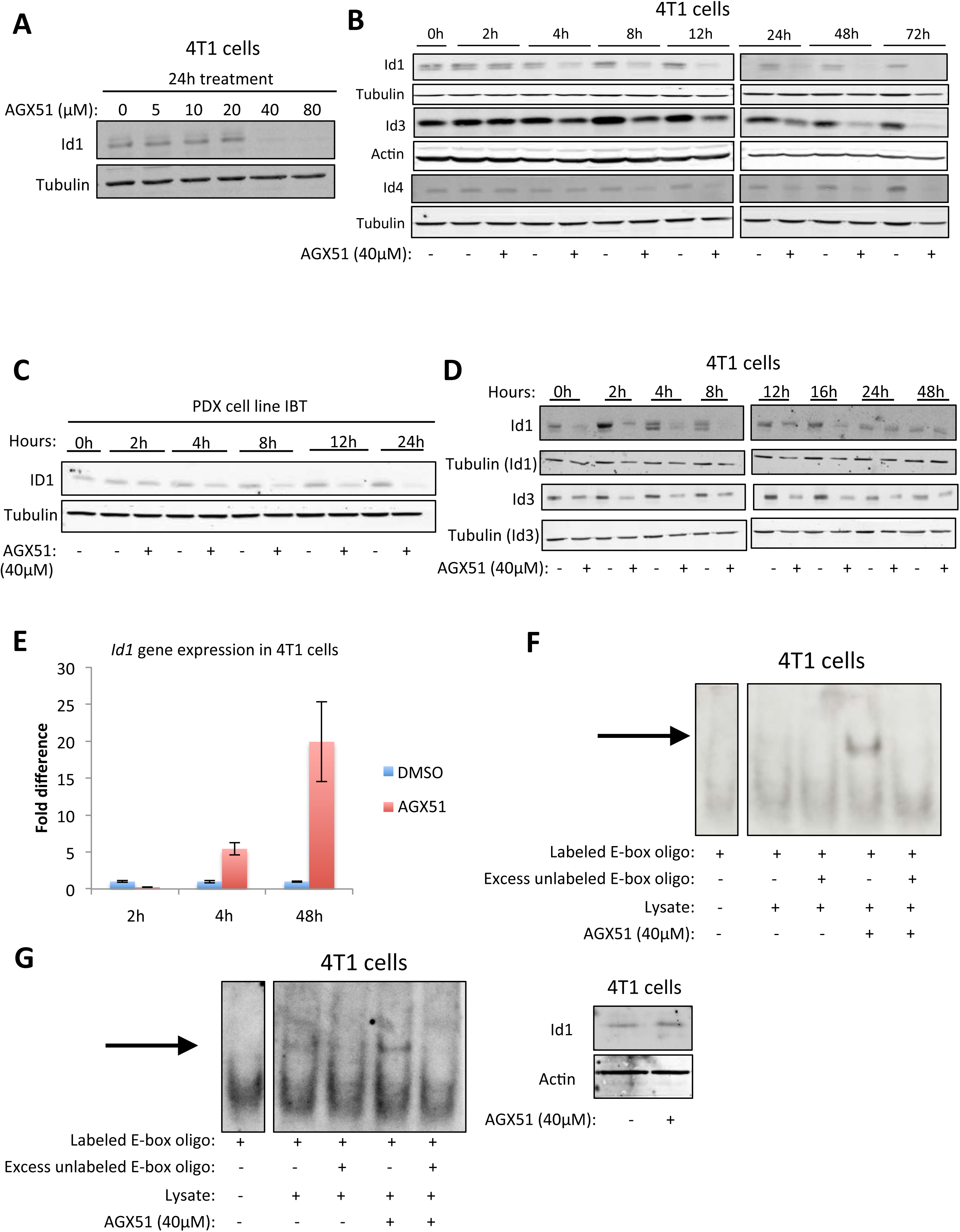
Effects of AGX51 on Id proteins. (A) Western blot for Id1 on whole cell lysates from 4T1 cells treated with 0-80 µM AGX51 for 24 hours. (B) Western blot for Id1, Id3 and Id4 on whole cell lysates from 4T1 cells treated with 40 µM AGX51 for 0-72 hours. Quantification of Western blot bands is shown in Figure S1A. (C) Western blot for ID1 on whole cell lysates from the PDX cell line IBT treated with 40 µM AGX51 for 0-24 hours. (D) Western blot for Id1 and Id3 on whole cell lysates from 4T1 cells treated with 40 µM AGX51 for 24 hours, after which compound was washed off and cells were collected for 0-48 hours post-compound removal. (E) qRT-PCR analysis for *Id1* in 4T1 cells treated with 40 µM AGX51 for 2, 4 or 48 hours. Mean fold difference with error bars showing SEM of technical triplicates. The experiment was performed at least three times. (F) EMSA on whole cell lysates from 4T1 cells treated with 40 µM AGX51 for 24 hours. Arrow indicates binding to DNA. (G) EMSA on whole cell lysates from 4T1 cells treated with 40 µM AGX51 for one hour, with corresponding western blot for Id1 shown to the right of the EMSA blot. Arrow indicates binding to DNA. Tubulin/Actin are used as protein loading controls. See also Figure S1.

To determine if the reduction in Id protein levels was secondary to reduced transcription, we measured *Id1* mRNA levels from 4T1 cells treated with AGX51 over a 48-hour period. By qRT-PCR, we observed a modest increase of *Id1* mRNA at early time points, with a dramatic increase in expression by 48 hours (5x and 20x, respectively, Figure 1E). A similar result was observed for *Id3* (data not shown). The increase in *Id1 and Id3* mRNA levels suggests that the cell is attempting to compensate for Id1 and Id3 protein loss, likely through the ability of the liberated E proteins to transactivate the *Id* promoters analogous to what is seen in *Drosophila melanogaster* (Bhattacharya and Baker, 2011) where daughterless (an E protein orthologue) activates *extramachrochaete* (an *Id* orthologue).

We next tested whether loss of Id proteins in response to AGX51 resulted in increased E protein-DNA binding. As expected, treating 4T1 cells with AGX51 for 24 hours resulted in an increase in E protein binding (Figure 1F). Similar EMSA results were obtained using cell lysates from HMLE *RAS Twist* cells, and the PDX cell line (Figure S1D). To determine if the loss of Id activity precedes Id protein loss, EMSAs were carried out after 1 hour of AGX51 treatment. An increase in E protein binding occurred at a time when no Id1 protein loss was observed (Figure 1G), suggesting that the increase in E protein binding is due to disruption of the Id1-E PPI as opposed to decreased overall Id levels. This conclusion is supported by direct evidence of the disruption of the ID1-E47 interaction observed in HCT116 cells (Wojnarowicz et al., 2019).

### Effects of AGX51 on cell growth

To determine the effects of AGX51-mediated Id protein loss on cell growth, we measured the IC50 of AGX51 in 4T1 cells and nine breast cancer cell lines representing the major breast cancer subtypes (ER+, HER2+, and TNBC), and three breast cancer PDX cell lines (Table S1). The PDX cell lines were the most sensitive to AGX51 treatment, displaying ∼2-fold lower IC50 values (∼15 μM) relative to the other cell lines (Table S1). Of the different breast cancer subtypes, the TNBC cell lines were most sensitive (∼25 μM) (Table S1). The IC50 of 4T1 cells, a model of TNBC (Aslakson and Miller, 1992; Pulaski and Ostrand-Rosenberg, 2001) was similar to that of the TNBC cell lines (Table S1). The effect of AGX51 on 4T1 cells was also determined using Alamar blue viability assays, which show a significant reduction (∼3-fold, p=0.0009) in viability after 24 hours of treatment with 40 μM AGX51 (Figure 2A).

**Figure 2.**
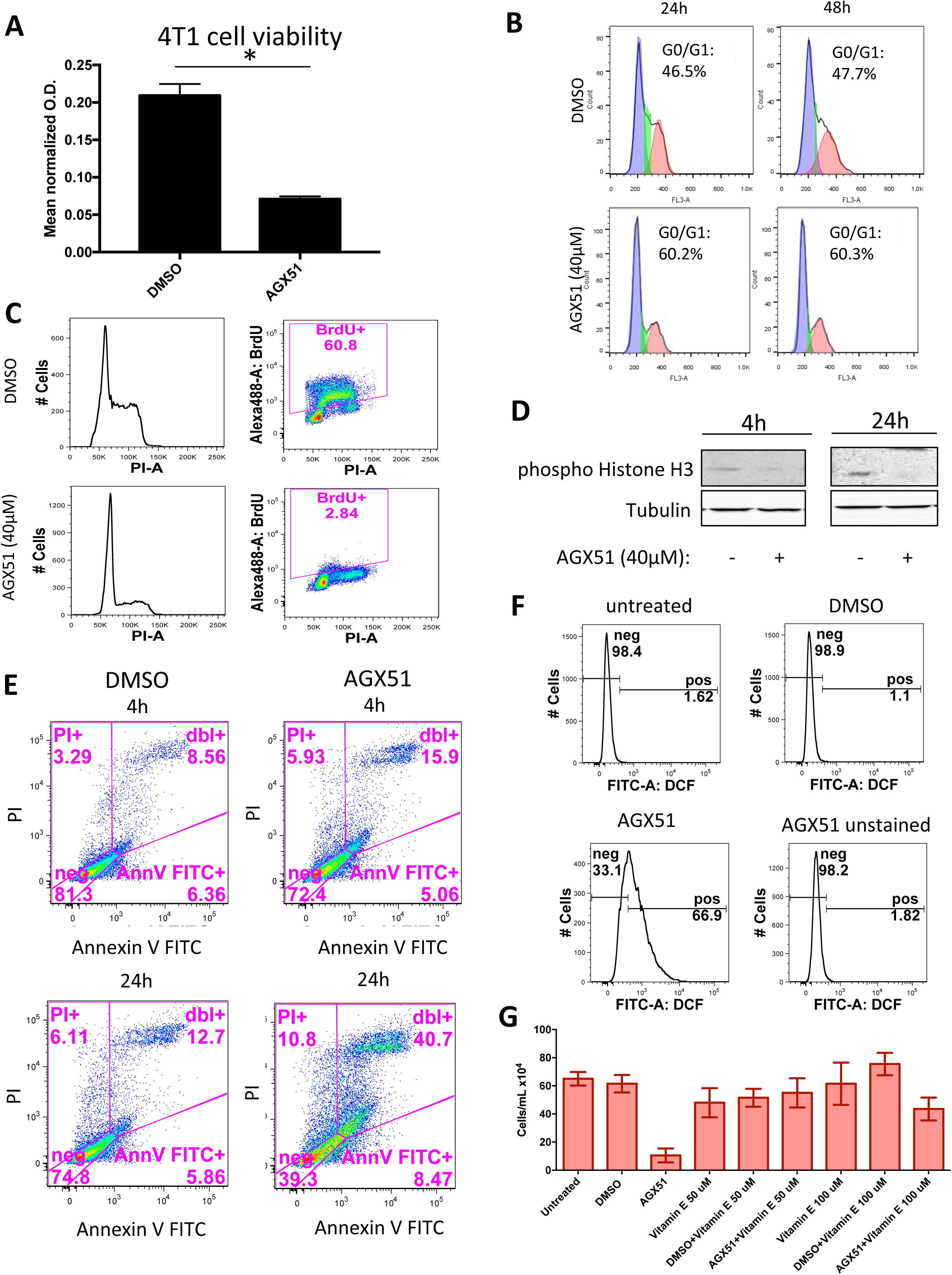
Effects of AGX51 on cells in culture. (A) Alamar blue cell viability assay on 4T1 cells treated with 40 µM AGX51 for 24 hours. Mean value of technical triplicates is plotted with error bars representing SEM. * Indicates p<0.001, as determined by unpaired t-test. The experiment was repeated at least three times. (B) Cell cycle analysis on 4T1 cells treated with 40 µM AGX51 for 24 and 48 hours; the experiment was repeated three times. (C) BrdU incorporation by 4T1 cells treated with 40 µM AGX51, or vehicle, for 24 hours as measured by flow cytometry. (D) Western blot for phospho-histone H3 on lysates from 4T1 cells treated with 40 µM AGX51, or DMSO for 4 and 24 hours. (E) Annexin V and PI staining of 4T1 cells treated with AGX51 for 4 and 24 hours, as analyzed by flow cytometry; the experiment was repeated two times. (F) Reactive oxygen species accumulation in 4T1 cells induced by AGX51 treatment, as determined by H2DCFDA staining followed by flow cytometry analysis. AGX51-treated cells not stained with H2DCFDA were analyzed as an auto-fluorescence control. The experiment was repeated three times. (G) Cell viability, as measured by trypan blue exclusion and cell counting of 4T1 cells treated with 40 µM AGX51 or DMSO for 24 hours, with or without 1 hour pretreatment with vitamin E (50 or 100 µM). See also Figure S2.

It has previously been reported that genetic loss of Id1 induces a G0/G1 growth arrest (Barone et al., 1994; Hara et al., 1994). After 24 and 48 hours of AGX51 treatment, 4T1 cells showed an accumulation in G0/G1 (Figure 2B). AGX51 treatment also led to a reduction in BrdU incorporation (Figure 2C) and phospho-Histone H3 levels (Figure 2D). After 24 hours of AGX51 treatment we saw an increase in the Annexin V+/PI+ fraction (12.7% vs. 40.7%), indicative of late apoptosis and necrosis (Figure 2E). We observed no overt increase in cleaved caspase 3 or cleaved PARP levels, suggesting that the mechanism of cell death is largely non-apoptotic (data not shown).

A feature of necrotic cell death is the production of reactive oxygen species (ROS). To determine the effect of AGX51 on ROS production in 4T1 cells, we treated cells with 40 μM AGX51 for 24 hours, stained with H2DCFDA and analyzed the cells by flow cytometry. We observed a significant increase in the fluorescein positive fraction upon AGX51 treatment, indicative of an increase in ROS production (Figure 2F). In fact, AGX51 increased ROS production to a greater degree than our positive control, H_2_O_2_ (Figure S2A). Similar effects were seen in HCT116 cells (Figure S2B). Furthermore, addition of 50 and 100 μM of the antioxidant vitamin E rescued cell viability, and reduced ROS levels in both cell lines (Figure 2G; Figure S2C-G). Vitamin E treatment also rescued the AGX51-mediated cell cycle arrest in HCT116 cells (Figure S2H), and 4T1 cells (data not shown) suggesting that the cell cycle arrest is secondary to ROS production. Interestingly, other free radical scavengers such as N-acetyl-cysteine (NAC), MnTBAP, MitoQ, MitoTEMPO, Apocynin, and Catalase did not rescue the AGX51-induced phenotypes (data not shown) indicating that an oxidized lipid may be a key mediator of AGX51 activity. Nonetheless, our findings are consistent with a recent report where siRNA-mediated Id1 and Id3 inhibition was associated with increased ROS levels (Bensellam et al., 2015) and indicate that a main mechanism of AGX51-mediated cell killing is ROS-mediated. Enforcement of ID1 expression has previously been reported to play an important role in circumventing Tgf-β’s tumor suppressive effect in pancreatic cancer. Depletion of the ID proteins by shRNAs has been shown to decrease the ability of pancreatic cancer cells to survive and form orthotopic tumors (Huang et al., 2019). We tested whether AGX51 would have similar effects on pancreatic cancer cells. In line with the results obtained in other cancer models that we tested, AGX51 caused a depletion of the ID1 and ID3 proteins in the pancreatic cancer cell line 806, starting at a dose between 4 and 20 µM (Figure 3A) with ID3 showing slightly less sensitivity as observed in other cellular and biophysical assays (Wojnarowicz et al., 2019). Furthermore, AGX51 decreased cell viability in a variety of pancreatic cancer cell and organoid lines, including mouse organoids (Figure 3B), mouse cell lines 806, NB44, and 4279 (Figure 3C), and human cell lines Panc1 and A21 (Figure 3D) with an IC50 of 5.5-19.5 µM. This corresponds to the range of concentration of AGX51 required for Id protein loss. Of note, when treated with 40 µM of AGX51, well-formed organoid structures collapsed (Figure 3E).

**Figure 3.**
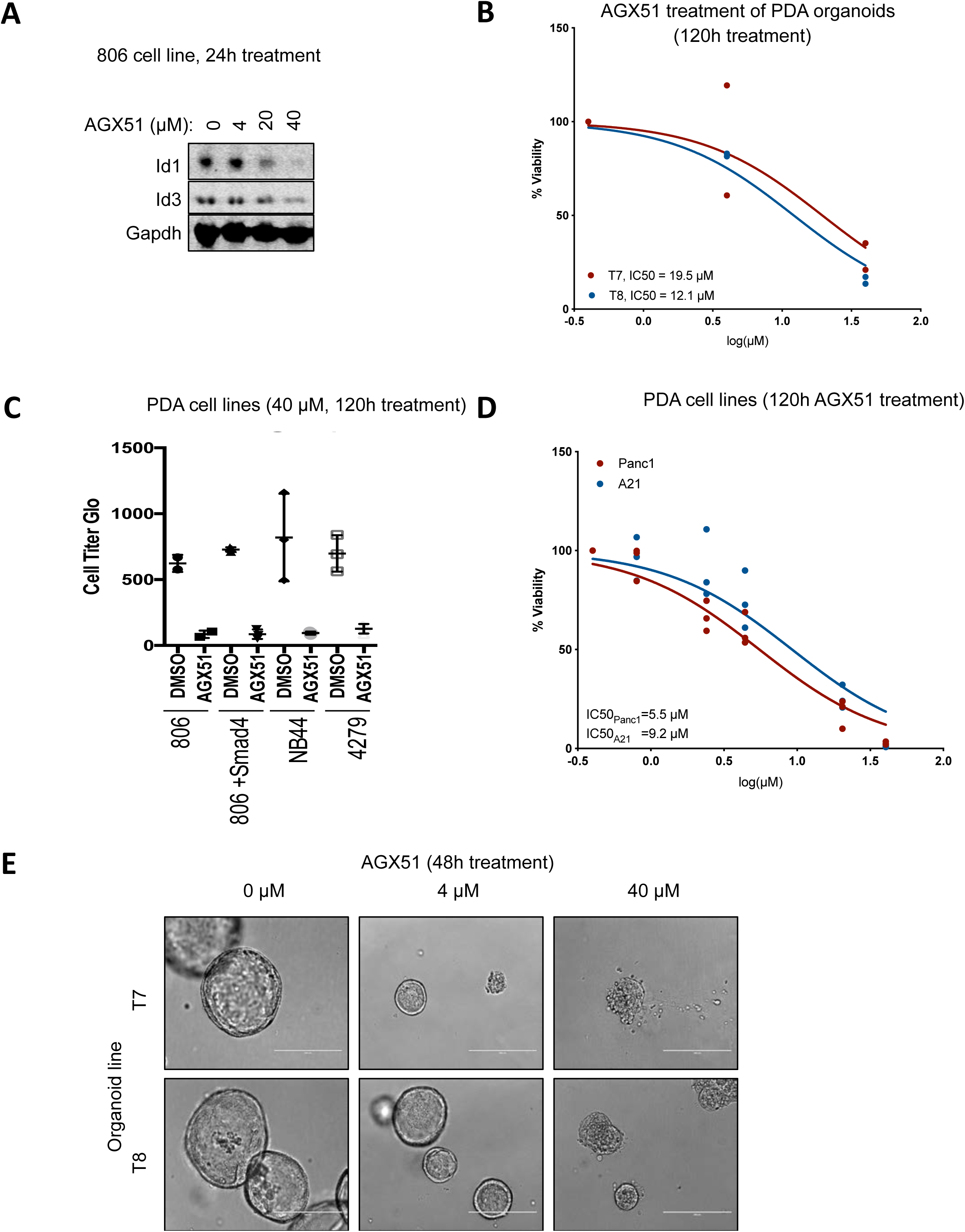
Effects of AGX51 on pancreatic ductal adenocarcinoma cells. (A) Western blot for Id1 and Id3 on whole cell lysates from 806 pancreatic ductal adenocarcinoma (PDA) cells treated with 4, 20 and 40 µM AGX51 for 24 hours. Gapdh used as loading control. (B) Dose response of PDA organoid lines T7 and T8 to AGX51 following 120 hours of treatment, with IC50 indicated. (C) Cell viability as assessed by Cell Titer Glo assay for PDA lines 806, 806+Smad4, NB44 and 4279 treated with 40 µM AGX51 or DMSO vehicle control for 120 hours. (D) Dose response of PDA cell lines Panc1 and A21 to 120 hours of AGX51 treatment, with IC50 indicated. (E) Bright field images of T7 and T8 PDA organoid lines treated with 4 and 40 µM AGX51 for 48 hours. Scale bars represent 200 µm.

### Specificity of AGX51 activity

A common concern with any novel small-molecule is specificity. Our results showing loss of Id proteins, an increase in liberated E proteins, and cell phenotypes consistent with genetic models of Id loss indicate that AGX51 is targeting Id proteins. However, AGX51 may also be affecting other, unintended targets. If Id proteins are the primary targets of AGX51 then we reasoned that reducing Id1 or Id3 levels by shRNA should increase the sensitivity of the cells to AGX51 by reducing the concentration of the target. A similar approach was used to show that reduction of Ras genes sensitized cells to a pan-Ras inhibitor (Welsch et al., 2017). Similarly, cells able to survive in the absence of Id proteins, such as quiescent cells should be resistant to AGX51. Upon partial knockdown of *Id1* or *Id3* by shRNAs (Figure S3A) the cells showed increased sensitivity to AGX51 (∼3-fold) (Table S1). 4T1 cells allowed to reach confluence under low serum conditions become quiescent and display near complete loss of Id expression (Figure 4A, B). Treating these cells with AGX51 led to a dramatic reduction in cell killing relative to cycling control cells. Together these data support the conclusion that the Id proteins are the primary targets in cycling cells.

**Figure 4.**
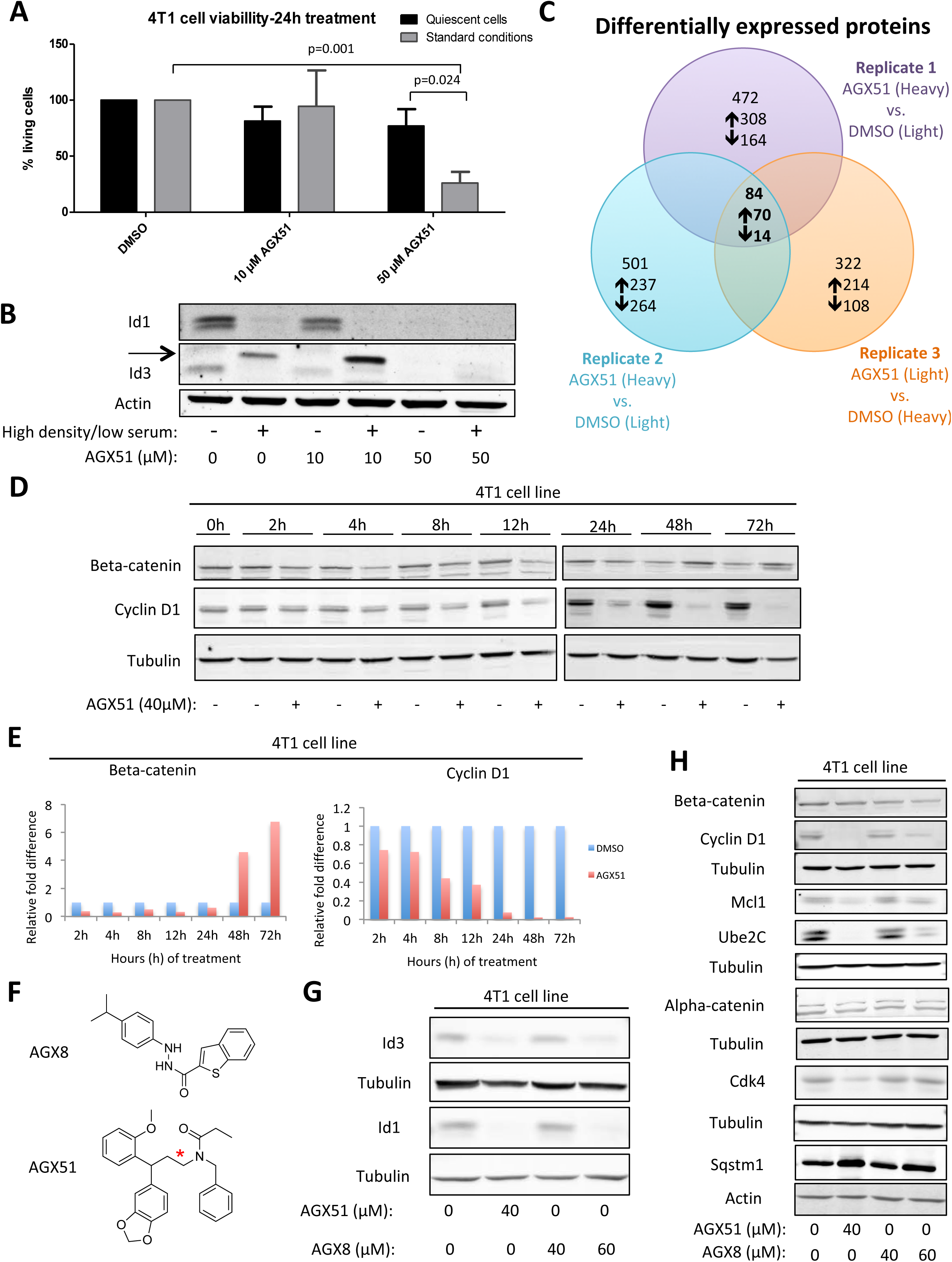
Assessing effect of AGX51 on quiescent cells and the whole proteome. (A) 4T1 cells were treated with AGX51 (10 or 50 µM), when they were cultured under standard conditions (standard density, standard serum (10%)) or under quiescence-inducing conditions (high density, low serum (1%)). Following 24 hours of AGX51 treatment, cell numbers were determined by cell counting and trypan blue exclusion. (B) Western blot for Id1 and Id3 on whole cell lysates from 4T1 cells presented in (A). Actin serves as the protein loading control. Arrow indicates non-specific band. (C) Venn diagram of differentially expressed proteins, as identified by mass spectrometry, from three replicates of SILAC-4T1 cells treated with 40µM AGX51 for four hours. (D) Western blot for Beta-catenin and Cyclin D1 on whole cell lysates from 4T1 cells treated with 40 µM AGX51 for 0-72 hours. (E) Quantification of Western blot shown in (D). (F) Structures of AGX8 and AGX51 (red asterisk indicating chiral center). (G) Western blot for Id3 and Id1 on whole cell lysates from 4T1 cells treated with AGX51 and AGX8 at the indicated concentrations. (H) Western blot for the indicated proteins on whole cell lysates from 4T1 cells treated with AGX51 and AGX8 at the indicated concentrations. Quantification of Western blot bands shown in (G) and (H) are in Figure S6. See also Figures S3-S6.

Having established that AGX51 disrupted PPI leading to proteasome-dependent degradation of ID proteins (Wojnarowicz et al., 2019), we wondered if levels of other proteins would be affected. To determine the full range of consequences of AGX51 exposure, we performed Stable Isotope Labeling of Amino acids in Cell culture (SILAC) on 4T1 cells treated with AGX51 or DMSO for four hours (at which point we would expect to see a decrease in Id1) and profiled the whole proteome by mass spectrometry. We identified 6869 proteins/protein clusters with an FDR <1%, where 2735 proteins were found in all three replicate experiments. As expected, Id1 was underexpressed in all three replicates (Fold differences: −1.39, −1.71, and −2.07). Using a 1.35-fold cut off, we identified 13 other underexpressed and 70 overexpressed proteins (Figure 4C, Table S2). Id2 and Id4 were not detected in this analysis and Id3 showed no significant change at four hours, consistent with the data shown above (see Figure 1B). The underexpressed proteins included Beta-catenin, Cyclin D1, Cdk4 and Ube2C, which are all involved in cell cycle progression and were validated by western blot (Figure 4D,E; Figure S3B). The underexpression of these proteins at early time points is modest by SILAC and hence quantitative differences by Western blot do not become prominent until later time points, and in some cases are transient (Beta-catenin). An additional three underexpressed proteins were also validated by western blot (Mcl1, Sqstm1 and Alpha-catenin) (Figure S3B) although the Sqstm1 inhibition was also transient. These changes were also observed in the PDX cell line and HMLE *RAS Twist* cells (Figure S4). We saw no change in *Beta-catenin* mRNA levels for 24 hours of AGX51 treatment of 4T1 cells, and only saw a significant decrease in *Cyclin D1* message at later time points (∼1-fold at 24 hours and ∼4-fold at 48 hours), suggesting that at early time points the changes in Beta-catenin and Cyclin D1 proteins may be occurring post-transcriptionally (Figure S5A). We tested the gene expression of *Cdk4* and *Mcl1*, and also saw no significant decrease in mRNA over time with AGX51 treatment, and saw upregulation at later time points (Figure S5B). Importantly, the loss of these proteins takes place prior to the changes in cell cycle distribution (Figure S5C), arguing against the possibility that these protein changes are secondary to a G0/G1 arrest.

Genetic suppression of *Id1-4* could test the possibility that the proteome changes seen after AGX51 treatment are secondary to Id protein loss but a timed, simultaneous reduction of all Id proteins prior to cell cycle arrest has been technically impossible. Instead we examined the effects of a structurally unrelated compound (AGX8) that also scored positive in the initial EMSA screen for perturbation of the Id1-E47 interaction (Wojnarowicz, et al, 2019). AGX51 and AGX8 contain 2 distinct chemical scaffolds with different interaction patterns. While AGX51 has only hydrogen acceptor groups, AGX8 has 2 hydrogen donors and 2 hydrogen acceptor groups. Treating 4T1 cells with AGX8 for 24 hours also led to reduced levels of Id1 and Id3 although the concentration needed to see this decrease was greater than that for AGX51 (60 vs. 40 μM) (Figure 4F,G; Figure S6A). In addition, the cell viability effects of AGX8 were less pronounced than those of AGX51 (Table S3). Western blot analyses found that the levels of Cyclin D1, Beta-catenin, Ube2C and Mcl1 decreased when 4T1 cells were treated with 60 μM AGX8, similar to 40 μM AGX51 treatment. Very modest decreases were observed for Cdk4 and Alpha-catenin with 40 μM AGX51 but were difficult to discern with 60 μM AGX8 (Figure 4H; Figure S6B). In addition, there was an increase in Sqstm1 expression after 24 hours of AGX8 treatment, similar to that seen with AGX51 treatment (Figure 4H, Figure S3B). The similarity in the effects of AGX51 and AGX8 on many of the proteins identified by SILAC is consistent with the idea that these changes are secondary to Id loss.

### Effects of AGX51 on tumor growth and lung colonization of cancer cells

#### Effect of AGX51 on MDA-MB-231 xenografts

In spontaneous primary tumors of the breast in the MMTV-Her2/neu model, genetic analyses indicated that loss of *Id1* and/or *Id3* was insufficient to inhibit tumor growth and needed to be combined with suboptimal doses of chemotherapy to cause regression (de Candia et al., 2003). We tested AGX51 in a primary tumor xenograft model using MDA-MB-231 cells with and without paclitaxel. Nude mice were implanted with MDA-MB-231 cells and tumors were allowed to grow to ∼100 mm^3^ before the initiation of the following treatments: vehicle (DMSO), paclitaxel 15 mg/kg qd (quaque die, i.e. once a day) on days 1-5, AGX51 60 mg/kg bid (bis in die, i.e. twice a day) for all 19 days, or paclitaxel 15 mg/kg on days 1-5 + AGX51 6.7 mg/kg, 20 mg/kg or 60 mg/kg bid for all 19 days. Also included was a treatment group that received 15mg/kg paclitaxel qd on days 1-5 and 60 mg/kg of AGX51 for just seven days. Paclitaxel treatment alone had a modest inhibitory effect on tumor growth while the addition of 6.7 and 20 mg/kg AGX51 to paclitaxel almost completely inhibited tumor growth and the 60 mg/kg dose resulted in tumor regression (Figure 5A, Table S4). Interestingly, just seven days of AGX51 treatment, in combination with paclitaxel, was sufficient to cause tumor regression, suggesting a long-lived anti-tumor effect that may persist even after treatment has stopped (Figure 5A, Table S4). AGX51 treatment alone did not suppress tumor growth. Thus, similar to the previously conducted genetic loss of function experiment (de Candia et al., 2003), AGX51 treatment leads to synergistic growth inhibition of primary breast tumors when combined with chemotherapy. While such synergy may be due to effects of Id loss on neoangiogenesis, in the current drug study we observed no obvious changes in angiogenesis or vascularization by gross observation of the tumors, histological evaluation or CD31 staining by IHC (data not shown, see also Discussion).

**Figure 5.**
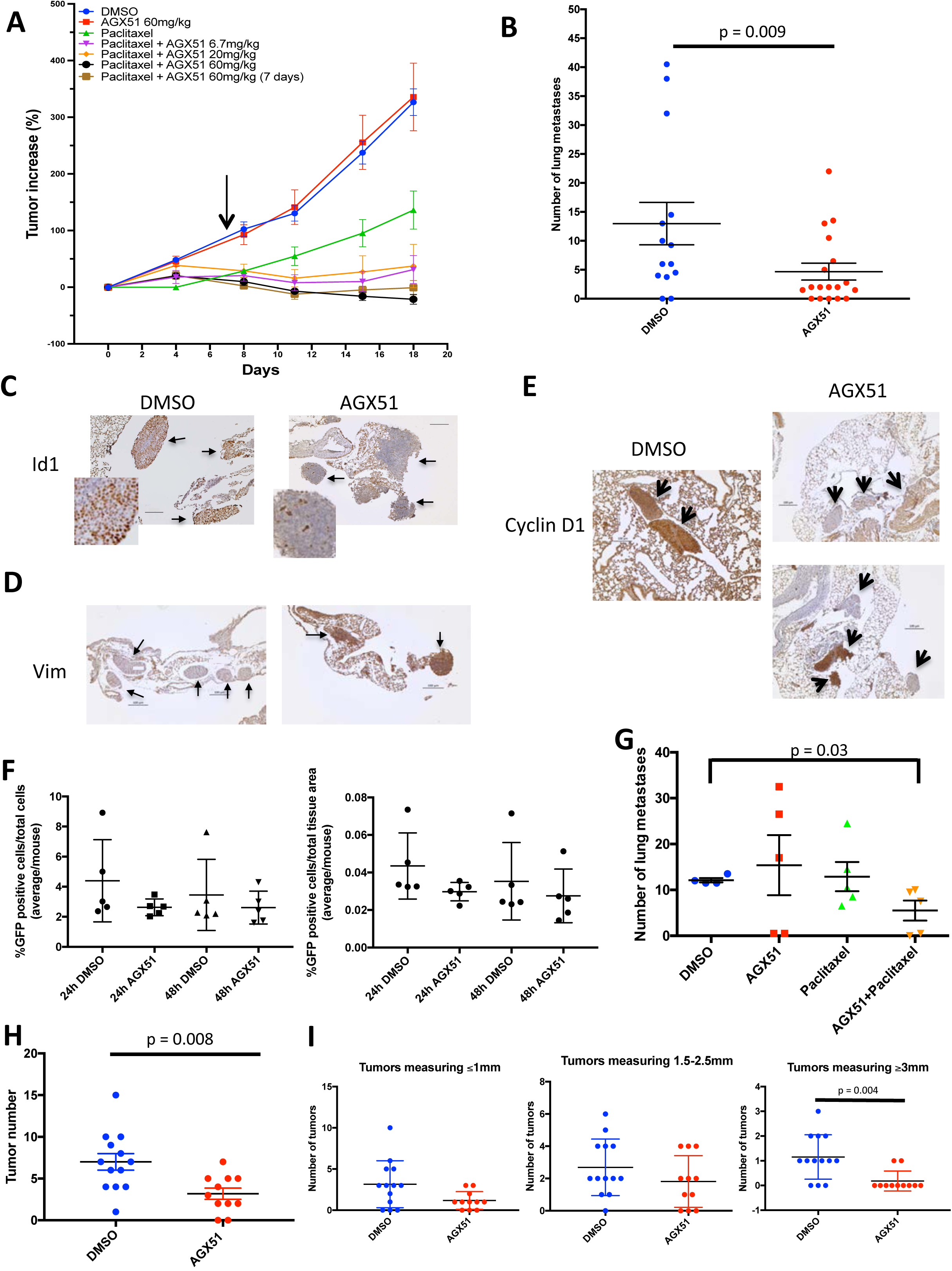
Effects of AGX51 on tumor growth and metastasis. (A) Mean tumor volume increase in athymic nu/nu mice implanted with MDA-MB-231 cells in the mammary fat pad and treated with AGX51, paclitaxel, AGX51+paclitaxel or vehicle control (n=5 mice per treatment group for all groups except paclitaxel alone where n=4 due to 1 death). Compared to controls, the effect of all AGX51+paclitaxel treatments was significant by weighted least squares (p≤0.001) and AGX51+paclitaxel treatments were also significantly different from AGX51 treatment alone (p≤0.001). The mean is plotted with error bars representing the SEM. (B) Average number of lung metastases in Balb/c mice injected with 5×10^4^ 4T1 cells via tail vein, treated with 50 mg/kg AGX51 bid or vehicle (n=18 or n=14, respectively). Means are plotted with error bars showing SEM. AGX51 treatment had a significant effect on lung tumor development (p<0.009 by weighted least squares). (C) Examples of Id1 IHC staining in lung sections from mice described in (B). (D) Examples of Vimentin IHC staining in lung sections from mice described in (B). (E) Examples of Cyclin D1 IHC staining in lung sections from mice described in (B), with examples of metastases from AGX51-treated mice showing negative (upper panel) and positive (lower panel) staining for Cyclin D1. Bars are 100 µm in length. (F) Quantification of the number of GFP-labeled 4T1 cells in the lungs of Ba l b /c mice injected with 5×10^4^ 4T1 cells via tail vein and treated with 50 mg/kg bid or vehicle for 24 and 48 hours (n=5 mice per treatment group). (G) Average number of lung metastases in Balb/c mice injected with 5×10^4^ 4T1 cells via tail vein and treated with AGX51, paclitaxel, AGX51+paclitaxel or vehicle once lung metastases were established (n=5 mice per treatment group). Means are plotted with error bars showing SEM. Compared to vehicle, AGX51+paclitaxel treatment had a significant effect on lung tumor growth (p<0.03 by weighted linear regression). (H) Number of colon tumors in AOM colon tumor model in A/J mice that were treated with DMSO (n=13) or AGX51 (11). AGX51 treatment resulted in a significant reduction in the number of tumors (p=0.008). (I) Size distribution of tumors in (H), grouped by treatment group (AGX51 or DMSO), where AGX51 treatment resulted in a significant reduction in tumors measuring ≥3 mm (p=0.004). Colon tumor count data was analyzed using the Wilcoxon Rank Sum test. For additional data pertaining to the effects of AGX51 on tumor growth and metastases see Figures S7-S9.

Of note, we have previously shown that after 14 days of daily treatment at the highest dose used in any of our studies (60 mg/kg bid), there were no statistically significant differences between the AGX51-treated animals and vehicle-treated animals with respect to weight, clinical chemistry and hematology plasma values, and histopathological evaluation of major organs at necropsy (Wojnarowicz et al., 2019). Similarly, when combining AGX51 bid with 15 mg/kg paclitaxel (qd for the first 5 days of a 14-day regimen), a regimen used in the xenograft studies described above, no overt toxicities were observed using any of the above listed parameters (Figure S7). Thus, in addition to showing anti-tumor efficacy, AGX51 is well tolerated in mice in this concentration range, alone and in combination with paclitaxel.

#### Effect of AGX51 on 4T1 cell colonization of the lungs

Genetic loss of *Id1/Id3* prevents the development of breast cancer lung metastasis by blocking the conversion of micro-metastases to macro-metastases (Stankic et al., 2013) and by blocking angiogenesis (Gao et al., 2008; Lyden et al., 1999). To determine whether AGX51 phenocopies these results, we used lung colonization assays, a model of experimental metastasis, where luciferase-labeled 4T1 cells, known to be metastatic, were injected into the tail veins of Balb/c mice and after 24 hours the mice were treated with 50 mg/kg of AGX51 i.p. bid or vehicle. Colonization of the lungs by 4T1 cells and their growth into tumors (referred to as metastases) was monitored weekly by *in vivo* bioluminescent imaging. The experiment was stopped four weeks following tail vein injection, due to the deteriorating condition of the control group. Bioluminescent imaging showed significant inhibition of lung metastasis development in the AGX51 group, and this correlated with the metastatic burden observed upon histopathological examination of lung tissues (p=0.009) (Figure 5B, Figure S8A). Treatment with AGX51 50 mg/kg once daily also decreased the number of lung metastases, and while still significant (p=0.0023) the effect was not as dramatic as that seen with the bid regimen (Figure S8B, C). While all macro-metastases showed strong positive IHC staining for Id1, in smaller lung metastases the Id1 staining was decreased in the AGX51 group, relative to metastases of a similar size in the DMSO group (Figure 5C). Conversely, as predicted from the genetic analyses (Stankic et al., 2013), Vimentin staining of small metastatic tumors in the AGX51 group was significantly higher than that in the DMSO group (Figure 5D), consistent with a reduction in *Id1/Id3* levels blocking the mesenchymal-to-epithelial transition during metastatic progression. Interestingly, Vimentin was one of the proteins found overexpressed in our SILAC analysis (see Table S2).

To characterize the effects of AGX51 on the expression of EMT-related proteins we assessed Vimentin, E-cadherin, Snail, Twist, and Zeb1 levels in AGX51-treated 4T1 cells. After an 8 hour exposure, we observed a decrease in E-cadherin and an increase in Vimentin but by 48 hours both proteins were increased. Decreases in Snail and Twist1 were observed throughout the time course, while Zeb1 levels were unchanged (Figure S9A). Thus AGX51 modulates the levels of some EMT-related proteins, however the precise mechanisms leading to these changes and their consequences remain to be elucidated. The changes observed do not appear to adhere to canonical EMT patterns, which may be evidence of AGX51 inducing partial EMT/MET. The occurrence of partial or intermediate EMT phenotypes in cancer is becoming increasingly recognized (Nieto et al., 2016; Pastushenko et al., 2018; Shibue and Weinberg, 2017). Finally, consistent with the reduced Cyclin D1 levels shown above, IHC staining for Cyclin D1 in the 4T1 lung metastases of the AGX51 group was significantly reduced relative to the DMSO group (p<0.00001) (Figure 5E). While all DMSO group metastases stained positive for Cyclin D1, about half of the AGX51 group was negative (Figure 5E upper panel of AGX51 group).

To determine whether AGX51 inhibits the extravasation and initial seeding at the secondary site or the progression of extravasated cancer cell outgrowth into tumors, we tail vein-injected mice with GFP-labeled 4T1 cells and treated the mice with AGX51 (50 mg/kg) or DMSO for 24 or 48 hours then stained the lungs for GFP (i.e. tumor cells) and quantified tumor cell number. There was no significant difference in the number of GFP positive cells between AGX51 and DMSO groups after either 24 or 48 hours of treatment (Figure 5F). These data are consistent with genetic models that found that *Id1/Id3* inhibition does not prevent extravasation of metastasizing breast cancer cells, but rather the conversion of micro- to macro-metastatic lesions (Gupta et al., 2007; Stankic et al., 2013).

To ascertain whether AGX51 is capable of inhibiting the growth of established lung tumors, we tail vein-injected 4T1 cells, as described above, and started treatment once lung signal was visible by bioluminescent imaging. Mice were then divided into four groups and treated with AGX51, paclitaxel, AGX51+paclitaxel or vehicle. Subsequent bioluminescent imaging and histopathological examination of the lungs found that while AGX51 or paclitaxel alone did not have an effect on the lung tumor growth, combination treatment significantly reduced the continued expansion of lung tumors relative to the other groups (p=0.03) (Figure 5G). In addition to suppressing the number of lung tumors, the tumors in the treated group were smaller in size (data not show). Thus, consistent with the primary tumor data, established lung tumors can be more effectively treated when AGX51 is combined with paclitaxel.

#### Effect of AGX51 on colon tumor development

To determine the effects of AGX51 in a sporadic tumor model, we turned to the azoxymethane (AOM) colon tumor model, a chemically induced autochthonous model of adenoma. Mice were administered AOM once a week for six weeks, followed by a three-week treatment break and then treated with AGX51 (15 mg/kg) or DMSO bid for three weeks. AGX51 treatment resulted in a significant decrease in the number of colon tumors (p=0.008). In addition, the tumors from AGX51 group were smaller than those in the DMSO group, and this difference was significant when tumors measuring >3 mm were compared (p=0.004) (Figure 5H, I). Thus AGX51 shows anti-tumor activity in a sporadic cancer setting.

### Investigating mechanisms of acquired resistance to AGX51

The efficacy of novel therapeutics can be hindered by cells developing resistance. Common mechanisms of acquired resistance include mutation of the drug binding site or overexpression of the target. As described above, mutation of the binding site would likely result in loss of Id1 activity, thereby blocking this mechanism of escape. Overexpression of the Id1 protein could potentially outcompete the drug. However, as described above, endogenous *Id1* mRNA levels increase 20-fold in response to AGX51 exposure but the protein is still degraded efficiently making this type of resistance unlikely. Nonetheless we tested the effects of AGX51 on HMLE *RAS Twist* cells lines overexpressing exogenous *ID1* mRNA (HMLE *RAS Twist ID1* cells). Western blot analysis revealed that despite significant ID1 overexpression, AGX51 treatment still dramatically reduced ID1 protein levels (Figure S10A). Similarly, 4T1 cells overexpressing exogenous *Id1* mRNA treated with AGX51 also showed a dramatic decrease in Id1 protein levels (Figure S10B). These data suggest that AGX51 is a potent Id1 degrader, even in the context of *Id1* mRNA overexpression.

To determine whether tumor cells could spontaneously acquire resistance to AGX51, we sought to identify subpopulations of cells capable of continued growth in the presence of AGX51. 4T1 cells were exposed to various AGX51 concentrations, with media replenished every 2-3 days. Above 25 μM no resistant colonies of cells were observed even after seven weeks of culture (∼3×107 cells tested). At 20 μM no overt cell death was observed, indicative of an extremely sharp dose response curve. We then decreased the concentrations to 24 μM and 22.5 μM, when Id loss after 24 hours is not as pronounced (Figure 6A). After six weeks in culture, we picked 30 seemingly viable clones but 29 eventually died after an average of ∼5 passages. One clone survived the 24 μM condition but grew at reduced rates relative to the starting population (Figure 6B), and eventually died after 11 passages.

**Figure 6.**
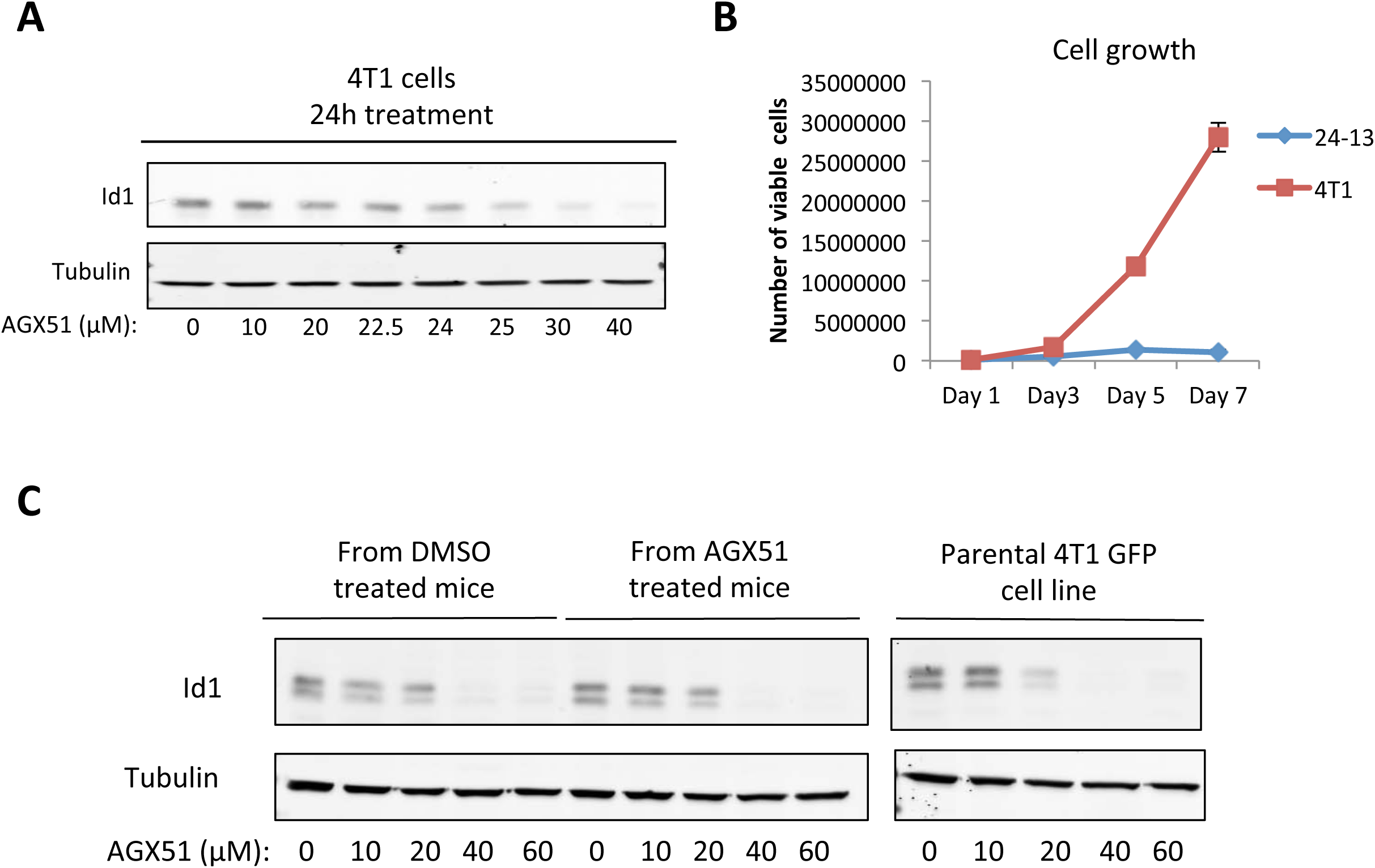
Generation of AGX51 resistant cell lines. (A) Western blot for Id1 on whole cell lysates from 4T1 cells treated with 0-40 µM AGX51 for 24 hours. (B) Cell growth of 4T1 cells and the AGX51 transiently resistant cell line clone 24-13, as determined by cell counting over seven days. Mean of technical replicates is shown with error bars representing SEM. (C) Cells were FACS isolated from the lungs of mice that were injected with 5×106 GFP-labeled 4T1 cells via tail vein and treated with 50 mg/kg bid or vehicle for two weeks. The isolated cells (from DMSO or AGX51-treated mice), and parental GFP-labeled 4T1 cells were treated with 0, 10, 20, 40 or 60 µM AGX51 for 24 hours and whole cell lysates were prepared. Western blot analysis for Id1 was carried out. Tubulin serves as a protein loading control. See also Figures S10 and S11.

None of the seven transiently resistant clones tested showed wild type levels of Id1 although about half did show a possible rebound in Id3 (Figure S11A, B). No Id2 expression was detectable in these cells. In 4/7 transiently resistant cell lines, there was no/low Id1 expression, while 3/7 transiently resistant cell lines maintained Id1 expression at the level of cells treated with AGX51 for 24 hours (Figure S11A). We also saw a decrease in Beta-catenin and Cyclin D1 levels in the transiently resistant cell lines (Figure S11C). Targeted sequencing of *Id1* and *Id3* coding regions in the three clones expressing detectable levels of Id1 and Id3 found no mutations, suggesting that mutation of the predicted compound-binding site, or other regions of the target proteins, is not the mechanism of transient resistance in these clones. While the growth rate of the clones cultured in 22.5 and 24 μM AGX51 was maintained for a time greater than that of the general population, this is not acquired resistance since all clones grew poorly and none remained viable after prolonged passaging. Furthermore, upon AGX51 removal, the cell growth rate and Id protein levels appeared to rebound to near untreated 4T1 levels, and fell again upon re-treatment with 22.5 or 24 μM AGX51 (not shown). No acquired resistance was similarly observed in four other cell lines tested: HMLE *RAS Twist*, SW480, HCT15 and HCT116 (data not shown). Thus, despite some cells being capable of propagation in 22.5 or 24 μM AGX51, the cells are under strong growth inhibitory pressure and fail to acquire genetic or epigenetic changes allowing for normal proliferation and Id expression in the presence of continued AGX51 exposure.

To determine if the small lung tumors that develop after AGX51 treatment are a result of acquired resistance *in vivo* we injected luciferase and GFP-labeled 4T1 cells into the tail veins of mice and 24 hours later started treating the mice with AGX51 (50 mg/kg, bid) or vehicle for two weeks. This primary lung metastasis model showed significant anti-tumor efficacy with AGX51 as a single agent, as described above. Consistent with previous results, we observed a significant reduction in lung tumor burden in the AGX51 group. After two weeks of treatment, we harvested and dissociated the lungs and collected the GFP positive cancer cells by FACS. After minimal passaging in cell culture (to obtain sufficient numbers of cells for western blot analysis), we re-challenged cells from both the AGX51 and DMSO treated animals, as well as the parental cell line, with AGX51. We saw no difference in AGX51 sensitivity between parental (no *in vivo* exposure), DMSO and AGX51 *in vivo* treated cells (Figure 6C). The IC50 values for AGX51 in the isolated cells were very similar between DMSO- and AGX51-treated mice (23.48 μM and 22.34 μM, respectively). This result demonstrates that the development of small lung tumors after AGX51 treatment is not due to acquired resistance inherent in the cell population but rather we suspect it is due to incomplete exposure of the tumor cells to AGX51.

## Discussion

In the present study, we characterized the effects of a novel small-molecule antagonist of Id proteins, AGX51, in cancer cell lines and mouse models of breast and colonic neoplasia. Importantly, AGX51 leads to a significant reduction of Id protein levels in all cell lines tested, lung tumors and ocular neovascularization (Wojnarowicz et al., 2019). Id protein destabilization after perturbation of its ability to antagonize E protein activity is consistent with genetic analyses in which coexpression of E proteins was shown to dramatically increase the stability of Id3 (Bounpheng et al., 1999; Deed et al., 1996). AGX51 appears to act as an Id protein antagonist and degrader, thereby accounting for its ability to phenocopy *Id* genetic loss of function phenotypes in multiple settings.

Several approaches were taken to address the question of AGX51 target specificity. SILAC analysis showed that after short exposure to AGX51 when the Id proteins first begin to get degraded, only 14/2735 other proteins were underexpressed. This limited effect on the proteome is unlikely to be a purely off-target effect as AGX8, a molecule structurally unrelated to AGX51 (see Figure 4F), also reduced Id protein levels and concomitantly the majority of the other proteins affected by AGX51. In addition, partial loss of *Id1* and *Id3* function reduced the IC50 of AGX51 (see Table S1) and quiescent cells in which Id proteins are absent are resistant to AGX51-mediated killing. These data support Id proteins being the primary targets of AGX51. Finally, the ability of AGX51 to recapitulate the effects of *Id* loss-of-function mutations in various settings also suggests that these proteins are the primary targets in cells: AGX51 induces a G0/G1 arrest; inhibits the conversion of micro-metastases to macro-metastases in the 4T1 breast cancer model of experimental lung metastasis; with paclitaxel leads to regressions of paclitaxel-resistant primary breast tumor xenografts; phenocopies the consequences of *Id1/Id3* loss in two models of pathologic neovascularization (Wojnarowicz et al., 2019); and decreases the viability of PDA organoids, as was seen with shRNAs to the Id proteins (Huang et al., 2019). AGX51 also showed anti-tumor activity in a sporadic model of colorectal neoplasia.

Despite repeated efforts, acquired resistance to AGX51 did not occur in cultured cells indicating that the escape frequency under our culturing conditions must be less than 1 in 10^7^ cells and perhaps significantly lower. These results suggest that despite the observed dramatic upregulation of *Id1* mRNA, Id protein levels are not restored efficiently nor are other pathways evoked, to compensate for Id loss. We also observed a lack of resistance in culture in cancer cells isolated from primary metastasis that persisted despite a two week AGX51 treatment regimen, suggesting that other factors such as drug exposure or the tumor microenvironment allowed for the survival of these cells. Another mechanism of resistance, mutating the binding site for the compound is also not observed since this likely results in an inactive protein given the highly conserved nature of this pocket and inability to tolerate directed mutations that maintain activity at these *loci in vitro* (Pesce and Benezra, 1993). Thus, by short-circuiting two major mechanisms of drug resistance (amplification of the Id target or mutation of the drug binding pocket) acquired resistance to AGX51 will be difficult to achieve and will probably serve to enhance the efficacy of AGX51 in a therapeutic setting.

The observed need for combining AGX51 treatment with chemotherapy to see anti-tumor efficacy in established mammary fat pad and lung tumors reflects the situation seen in genetic analyses where Her2/neu breast tumor regression required chemotherapy in Id knockout mice (de Candia et al., 2003). The reasons for this are not yet understood but it may be due to there being very few Id positive resting stem cells, as has been observed in other models (Cook et al., 2016), requiring combination treatment to hit two tumor populations (Id positive and Id negative) to see significant effects. Future studies will be required to resolve this question.

Id protein expression in normal adult tissue stem cells is required in response to various stresses: in colonic stem cells, Id1 is required to minimize inflammation after DSS induced chemical injury in a model of colitis (Zhang et al., 2014); in long-term repopulating hematopoietic stem cells they are required for optimal cellular expansion after ablation of the progenitor pool or to maintain long term serial transplantation (Jankovic et al., 2007; Perry et al., 2007); in neural stem cells, Id1 and Id3 are expressed in quiescent stem cell populations but show pronounced phenotypes only when those cells are called into cell cycle upon AraC ablation of progenitor cells (Nam and Benezra, 2009); Id proteins are expressed at very low levels in the liver but are reactivated during regeneration in response to injury (Le Jossic et al., 1994); Id1 is not expressed in normal human epidermis but may be a key regulator of epidermal wound healing (Schaefer et al., 2001). These observations likely account for the low toxicity of AGX51 in animals but suggest caution in the use under conditions of duress. In cancer, a similar phenomenon may be taking place since in glioma, Id1 and Id3 appear to be in a resting stem-like cell which resists killing of cycling cells by ionizing radiation (Cook et al., 2016). We have recently observed Id1 associated with a resting state in cholangiocarcinoma and triple negative breast cancer (R.B. unpublished). It will be of great interest to determine if combining standard cytotoxic therapies and AGX51 would kill the Id1-negative cycling population and the Id1-positive stem-like cells called into cycle, respectively.

Importantly, other PPIs have been shown to be critical for Id activity in the stem cell character of other tumors: Id2 for example facilitates stemness in glioma by its association with the VHL complex and antagonism of the ubiquitin mediated proteolysis of Hif2α (Lee et al., 2016), so determining the effects of AGX51 on this pathway will be of great interest. Given the complete suppression of gliomagenesis upon loss of *Id1*, *Id2* and *Id3* in a murine model (Niola et al., 2013) and the ability of AGX51 to antagonize all members of the family, it is likely that sufficient exposure of these brain tumors to AGX51 will provide some therapeutic benefit.

In conclusion, the first-in-class small-molecule antagonist of the Id protein family, AGX51, dramatically reduces Id protein levels and cancer cell viability, as well as inhibits tumor growth. The ability of AGX51 to phenocopy the consequences of *Id* genetic loss suggest that in addition to being a useful biologic tool, AGX51 can serve as a lead compound that can be developed to provide clinical benefit for a variety of Id-related cancers.

## Acknowledgements

The authors are grateful to members of the Benezra Lab, past and present, for helpful discussions throughout the course of this work. We are also grateful to Andrea Rizzi for additional supportive modeling work of AGX51 interactions. We thank the MSKCC Molecular Cytology Core Facility for assistance in image analyses and the MSKCC Flow Cytometry Core Facility for assistance with flow cytometry. RB gratefully acknowledges support from the Sloan Kettering Institute and NIH grant P30 CA008748. RB is also supported by grants from the NCI (PO1 CA094060) and The Breast Cancer Research Foundation. AJD is supported by funding from the Breast Cancer Research Foundation and the Botwinick-Wolfensohn foundation in memory of Mr. and Mrs. Benjamin Botwinick. Work was partially supported by 1R43CA150448 01 from NCI/NIH to WAG and National Institute of Health grants to AI (R01CA101644 and R01CA131126). PMW is a recipient of a Postdoctoral Training Award from the Fonds de recherche du Quebec – Sante (FRQS). MGE was supported by fellowships issued by the Valencian Government of Spain (GVA) and the European Social Fund (ACIF/2018/004 and BEFPI/2018/029).

## Author contributions

Conceptualization, R.B; Methodology, P.M.W., R.K.S., J.P., D.C.M., and R.B.; Formal Analysis, P.M.W., R.K.S., J.P., D.C.M., and X.K.Z.; Investigation, P.M.W., M.G.E., B.D., Y.C., R.S., S.X., R.K.S., J.P., D.C.M.; Resources, J.H.H., and V.K.; Writing – Original Draft, P.M.W., and R.B.; Writing – Review & Editing, R.B., P.M.W., D.C.M., A.D. A.J.D., and M.G.E.; Visualization, P.M.W., and R.B.; Supervision, O.O., L.N., N.R., A.I., A.L., and R.B.; Funding Acquisition, A.I., A.J.D., and R.B.

## Methods

### Electrophoretic mobility shift assays

Electrophoretic mobility shift assays (EMSAs) were carried out on whole cell lysates from AGX51-treated cells and the EMSA was performed as described previously (Tournay and Benezra, 1996).

### Cell lines

The 4T1 murine mammary tumor cell line, MDA-MB-157, MDA-MB-436, MDA-MB-231, MDA-MB-453, MDA-MB-361, BT-474, SK-BR-3, MCF-7, T47-D and HCT116 were purchased from ATCC (Manassas, VA, USA) and grown in RPMI or DMEM (Dulbecco’s Modified Eagle Medium) media supplemented with 10% FBS (fetal bovine serum), 1% penicillin-streptomycin and 1% L-Glutamine. MDA-MB-231 cells used in xenograft studies were obtained from NCI NIH (Bethesda, MD). Luciferase labeled 4T1 cells were described previously (Granot et al., 2011), as were luciferase and GFP-labeled 4T1 cells (Zimel et al., 2017). 4T1 cells overexpression Id1 were derived by transducing cell with pBabe-*Id1* plasmids as described previously (Stankic et al., 2013). HMLE *RAS Twist* and HMLE *RAS Twist ID1* cells were described previously (Stankic et al., 2013) and grown in MEBM (Mammary Epithelial Cell Growth Medium) (Lonza), supplemented with BPE (Bovine Pituitary Extract) (70 μg/mL), hEGF (human Epidermal Growth Factor) (5 ng/mL), hydrocortisone (0.5 μg/mL), insulin (5 μg/mL), GA-1000 (Gentamicin 30 μg/mL and Amphotericin 15 ng/mL), hTransferrin (5 μg/mL), and isoproterenol (6 μM). 4T1 cells were authenticated by short tandem repeat analysis and karyotyping. Other cell lines have not been authenticated since being obtained from their source. The cell lines were not tested for mycoplasma contamination recently but routine testing had been negative for years prior and no growth defects associated with such contamination have been observed in any of the cell lines used in the studies presented.

### Patient derived xenograft cell lines BR7, BR11, and IBT

PDXs were established directly from breast cancer bone metastases specimens surgically resected from patients with informed consent (IRB protocol #97-094). BR7 and BR11 specimens are obtained from ER positive (HER2 negative and PR negative) metastatic breast cancer patients, while the IBT specimen was obtained from a metastatic, triple negative breast cancer patient. Fresh tumor tissues were quickly washed with ice cold PBS and minced into about 1 mm3 pieces in MEM medium (without FBS) using sterile razor blades. A fraction of the minced original tumor tissues was incubated with Collagenase/Hyaluronidase enzyme mix (1,000 Units, Voden Medical) in MEM medium without FBS (5 mL/250 mg tissue) for 2-4 hours. The dissociated tumor tissues were then filtered throughout a 70 μm nylon filter, cells were concentrated by centrifugation at room temperature and seeded to derive respective primary cell cultures in MEM medium with 3% FBS (Sigma). The primary cell cultures were transduced with the fluorescent td tomato-/EGFP-luciferase fusion protein expressing lentiviral vectors for 18-24 hours, the primary cells were then maintained in MEM media supplemented with 3% FBS, 1% penicillin-streptomycin and 1% L-Glutamine. Aliquots of the primary cell cultures were also cryopreserved following a minimal number (3-4) of *in vitro* passages.

### Pancreatic ductal adenocarcinoma cells and organoids

The human pancreatic cancer cell line Panc1 was obtained from ATCC in 2012, and A21 was obtained in 2016 from Dr. Christine Iacobuzio-Donahue (Jones et al., 2008). Mouse pancreatic cancer lines 806 (KrasG12D; Ink4a-/-; Smad4-/-) and NB44 (KrasG12D; Ink4a-/-) were obtained from Dr. Nabeel Bardeesy, and 4279 (KrasG12D; Ink4a-/-) was generated in Dr. Joan Massague’s laboratory. Mouse pancreatic organoid cell lines T7 and T8 were obtained from Dr. David Tuveson (Boj et al., 2015). Pancreatic spheroids were grown in Ultra Low Attachment Culture plates (Corning) in DMEM supplemented with Glutamax (2 mM) and heparin (5 µg/mL). Pancreatic organoids were embedded in Matrigel with Advanced DMEM/F12 (Gibco, 12634-028) supplemented with B-27 (Life Technologies, 12587-010), HEPES (10 mM), 50% Wnt/R-spondin/Noggin-conditioned medium (ATCC, CRL-3276), Glutamax (Invitrogen, 2 mM), N-acetyl-cysteine (Sigma, 1 mM), nicotinamide (Sigma, 10 mM), epidermal growth factor (Peprotech, 50 ng/mL), gastrin (Sigma, 10 nM), fibroblast growth factor-10 (Peprotech, 100 ng/mL), A83-01 (Tocris, 0.5 µM) as previously described (Boj et al., 2015). All cell lines and organoids were maintained at 37°C and 5% CO2. Cell viability was measured using Cell Titer-Glo (Promega) according to manufacturer’s instructions.

### Generating Id knockdown cell lines

To generate 4T1 cells containing inducible short hairpins against *Id1* or *Id3*, short hairpins against *Id1* (TGCTGTTGACAGTGAGCGAGCGTGTTTCTGTTTTATTGAATAGTGAAGCCA CAGATGTATTCAATAAAACAGAAACACGCGTGCCTACTGCCTCGGA) and *Id3* (TGCTGTTGACAGTGAGCGCGCCCTGATTATGAACTCTATATAGTGAAGCCA CAGATGTATATAGAGTTCATAATCAGGGCATGCCTACTGCCTCGGA) were cloned into the tet-regulated all in one retroviral miR-E expression vector RT3REVIR (Fellmann et al., 2013). Retroviral particles were produced by lipofectamine-mediated transfection into 293 GP2 cells. Forty-eight hours after transfection, viral-containing media was collected, filtered and added to 4T1 cells in the presence of 8 μg/mL polybrene. Transduced cells were selected for Venus positivity by FACS. To induce short hairpin expression cell culture media was supplemented with doxycycline (5 μg/mL). RT3REVIR expressing a short hairpin against Renilla was used as a control (shCTL) in experiments.

### Immunoblotting

For immunoblotting, cells were collected by trypsinization, washed with PBS and lysed in homogenization buffer (0.3 M sucrose, 10 mM Tris (pH 8.0), 400 mM sodium chloride, 3 mM magnesium chloride, 0.5% NP40/IGEPAL, 100 μg/mL Aprotinin+Protease inhibitor cocktail (Roche # 11 836 153 001). Proteins were separated by SDS-PAGE, transferred to a membrane (LI-COR), probed with primary antibodies overnight at 4oC, and probed with secondary antibodies (LI-COR) for 1-2 hours at room temperature. Proteins were visualized using the LI-COR Odyssey Infrared Imaging detection system. The following primary antibodies were used Id1, Id2, Id3, Id4 (195-14, 9-2-8, 17-3, 82-12, respectively, all from Biocheck), phospho Histone H3 (9701, Cell Signaling), Cyclin D1 (2978, Cell Signaling), Beta-catenin (9562, Cell Signaling), Cdk4 (sc-260, Santa Cruz), Mcl1 (5453, Cell Signaling), Ube2C (14234, Cell Signaling), Sqstm1 (H00008878-M01, Abnova), Alpha-catenin (C2081, SIGMA-ALDRICH), E-cadherin (3195 Cell Signaling), Vimentin (5741, Cell Signaling), Snail (3879, Cell Signaling), Twist (ab50887, Abcam), Zeb1 (NBP1-05987, Novus Biologicals), Actin (A2066, SIGMA), Tubulin (T4026, SIGMA). Western blot quantification was carried out using channel 700 and channel 800 intensity data from Odyssey application software version 3.0.30 (LI-COR), subtracting blank values and normalizing to tubulin.

### qRT-PCR

RNA was extracted using the RNeasy kit (Qiagen, Valencia, CA, USA) and cDNA was generated from 1 μg of RNA using SuperScript IV First-Strand Synthesis System (Invitrogen, Grand Island, NY, USA). Quantitative PCR was performed using SYBR Green QuantiTect Primer Assay (Qiagen) according to manufacturer’s instructions in a 7900HT Fast-Real Time PCR System Instrument (Applied Biosystems, Grand Island, NY, USA). Primer pairs for the individual genes were obtained from the bioinformatically validated QuantiTect Library and are as follows: *Id1* (QT01743756), *Actb* (QT00095242), *Ctnnb1* (QT00160958), *Ccnd1* (QT00154595), *Cdk4* (QT00103292), *Mcl1* (QT00107436), *Idh2* (QT00147301), *Idh3b* (QT00117341), *Nbas* (QT00280483), *Ifrd1* (QT00093401), *Atp5h* (QT00175070), *C1qbp* (QT01057777), *Cox5a* (QT00164234), *Dlst* (QT00163275), *Glud1* (QT00103411), *Vim* (QT00159690). The fold changes in gene expression were calculated using the delta-delta CT method.

### Cell viability assays

Alamar blue cell viability assay (Invitrogen) was carried out according to manufacturer’s instructions. Briefly, 5000 4T1 cells were seeded in a 96-well plate, the next day they were treated with 40 μM AGX51 for 24 hours, after which point a 1:10 dilution of Alamar blue cell viability reagent was added to the cells and absorbance was measured 2, 3, 4, 5 and 6 hours after addition of the reagent using a plate reader (Synergy 2, BioTek).

For IC50 determination, cancer cell lines were seeded in a 96 well plate (5000 cells per well). After overnight incubation, cells were treated with AGX51 (0, 5, 10, 20, 40, 60 μM) and incubated for 24, 48 and 72 hours, each condition was done in triplicate. At each time point 40 µL of MTT reagent (5 mg/mL) was added per well and the cells were incubated for four hours. Following incubation, media was aspirated and 200 µL DMSO was added per well. Absorbance was then measured at 570 nm using a plate reader (Synergy 2, BioTek).

To determine whether vitamin E could rescue AGX51 cell viability effects, cells were pretreated with vitamin E ((±)-α-Tocopherol, SIGMA) (50 or 100 µM) for 1 hour. Following the 1 hour treatment, cells were treated with 40 µM AGX51/DMSO for 24 hours. Cell viability was then determined by trypan blue exclusion and cell counting.

Cell proliferation of the AGX51 resistant clone (24-13) was determined by plating 60,000 24-13 or 4T1 cells and collecting and counting the cells on days 1, 3, 5, and 7 after plating using a hemocytometer with the addition of trypan blue to exclude dead cells.

### Cell cycle analysis

Cells were treated with AGX51 or DMSO, or vitamin E (as described above), collected by trypsinizaton, washed with 1X PBS, resuspended in 500 μL 1X PBS and then diluted with 6 mL 70% ethanol and stored at -20°C until analysis. For cell cycle analysis cells were centrifuged 1000 rpm for 5 minutes, washed with 1X PBS and then resuspended in 0.5 mL PI/RNase staining buffer (550825, BD Biosciences), incubated for 15 minutes at room temperature and analyzed by flow cytometry (LSR II).

AGX51 or DMSO treated cells were stained with 10 μM BrdU reagent (550891, BD Biosciences) for one hour. The cells were then collected, resuspended in 70% ethanol, washed with 1X PBS, incubated with 2 M HCl, washed in 1X PBS and 1X PBS-Tween-BSA, and then incubated with an Alexa Fluor 488 anti-BrdU antibody (558599, BD Biosciences) for 15 minutes and finally resuspended in RNase/PI buffer (550825, BD Biosciences) and analyzed by flow cytometry.

### Annexin/PI staining

To detect apoptosis and necrosis we used the BioVision Annexin V-FITC Apoptosis detection kit according to the manufacturer’s instructions. Briefly, 4T1 cells were treated with 40 μM AGX51, typsinized and resuspended in 1X Binding Buffer to which 5 μL Annexin V-FITC and 5 μL propidium iodide (PI) (50 μg/mL) were added. The cells were incubated for 5 minutes protected from light and then analyzed by flow cytometry (LSR II).

### Measuring ROS

4T1 cells were treated with 40 μM AGX51 for 24 hours, and then 10 μM 2’,7’-dichlorodihydrofluorescein diacetate (H2DCFDA) (Life Technologies) was added to the cells and they were incubated for one hour. Cells were washed twice with PBS, trypsinized and resuspended in PBS-5%FBS. H2DCFDA is converted to fluorescent DCF upon oxidization, and serves as a readout for ROS generation. Fluorescence was measured using flow cytometry (LSR II). As a positive control for ROS production, cells were treated with 100 μM of H_2_O_2_.

### Testing the effects of AGX51 on quiescent cells

To induce a senescent state prior to AGX51 treatment, 4T1 cells were plated to reach high density at the time of AGX51 treatment. The day after plating the cells, media was changed to low serum media (1% FBS). After 24 hours in low serum media, the cells were treated with AGX51. Cells plated at standard density and maintained in 10% FBS served as controls. Following the 24 hours of AGX51 treatment, cell numbers were determined by cell counting and trypan blue exclusion and cells collected for whole cell lysate extraction for western blot analyses.

### Generation of SILAC cell lines, and in-gel digestion for mass spectrometry

4T1 cells were grown in DME media supplemented with 10% FBS and penicillin and streptomycin and either unlabeled L-arginine (Arg0) and L-lysine (Lys0) at 50 mg/L or equimolar amounts of the isotopic variants [U-13C6,15N4]-L-arginine (Arg10) and [U-13C6]-L–lysine (Lys6), (Cambridge Isotope Laboratories). After five cell doublings, cells were >99% labeled with the isotopes, as determined by mass spectrometry. Cell lysates were prepared as described above. 1 mg of total protein from each light and heavy labeled cell line was mixed and the total 2 mg of protein was fractionated using 4-12% gradient SDS-PAGE. Each gel lane cut in equal pieces and gel slices were washed with 1:1 (Acetonitrile: 100 mM ammonium bicarbonate) for 30 minutes. Gel slices were then dehydrated with acetonitrile for 10 minutes until gel slices shrunk and excess acetonitrile was removed before slices were dried in speed-vac for 10 minutes without heat. Gel slices were reduced with 5 mM DTT for 30 minutes at 56oC in an air thermostat, then chilled to room temperature, and alkylated with 11 mM IAA for 30 minutes in the dark. Gel slices were washed with 100 mM ammonium bicarbonate and acetonitrile for 10 minutes each. Excess acetonitrile was removed and dried in a speed-vac for 10 minutes without heat and gel slices were rehydrated in a solution of 25 ng/μL trypsin in 50 mM ammonium bicarbonate on ice for 30 minutes. Digestions were performed overnight at 37°C in an air thermostat. Digested peptides were collected and further extracted from gel slices in extraction buffer (1:2 vol/vol) 5% formic acid/ 50% acetonitrile) at high speed shaking in an air thermostat. Supernatant from both extractions were combined and dried down in a vacuum centrifuge. Peptides were desalted with C18 resin-packed stage-tips. Desalted peptides were lyophilized and stored at −80°C until further use.

### LC-MS/MS and whole proteome analysis

Desalted peptides were dissolved in 3% acetonitrile/0.1% formic acid and were injected onto a C18 capillary column on a nano ACQUITY UPLC system (Waters) that was coupled to the Q Exactive plus mass spectrometer (Thermo Scientific). Peptides were eluted with a non-linear 200 minute gradient of 2-35% buffer B (0.1% (v/v) formic acid, 100% acetonitrile) at a flow rate of 300 nL/min. After each gradient, the column was washed with 90% buffer B for 5 minutes and re-equilibrated with 98% buffer A (0.1% formic acid, 100% HPLC-grade water) for four minutes. MS data were acquired with an automatic switch between a full scan and 10 scan data-dependent MS/MS scan (TopN method). Target value for the full scan MS spectra was 3 X 106 charges in the 380-1800 m/z range with a maximum injection time of 30 ms and resolution of 70,000 at 200 m/z.in profile mode. Isolation of precursors was performed with 1.5 m/z. Precursors were fragmented by higher-energy C-trap dissociation (HCD) with a normalized collision energy of 27 eV. MS/MS scans were acquired at a resolution of 17,500 at 200 m/z with an ion target value of 5×104 maximum injection time of 60 ms and dynamic exclusion for 15 s in centroid mode. A total of 6346 proteins (5604 clusters) with an FDR <1% were identified across the three independent pairs of samples analyzed (Four-hour treatment of 4T1 cells with AGX51 or DMSO vehicle control). The data was analyzed using Scaffold Q+ version 4.4.5 (Proteome Software, Portland, Oregon, USA). Briefly, fold differences of the fold change ratios were calculated for each pair of samples (AGX51 vs. DMSO) and proteins with fold differences (at least 1.35-fold) in the same direction of change (overexpressed or underexpressed) in each of the three pairs were identified (see Table S2).

### Animal studies

Animal studies were carried out in accordance with institutional regulations (IACUC protocol 06-10-025) in a non-blinded fashion. Weight measurements and standard blood analyses were carried out on 8-12 week-old male CD1 mice (n=5 per treatment group) dosed q5d with DMSO, 15 mg/kg qd paclitaxel, 60 mg/kg AGX51 bid, or a combination of paclitaxel and AGX51. Blood samples were collected by retro-orbital puncture on day six, 12 hours after the last dose. Plasma was analyzed for clinical chemistry (albumin, alkaline phosphatase, alanine aminotransferase, amylase, total bilirubin, blood urea nitrogen, calcium, phosphorus, creatine, glucose, sodium, potassium, total protein and globulin) and hematology (white blood cell, lymphocyte, monocyte, granulocyte, red blood cell, hemoglobin, hematocrit and platelet) concentrations.

Orthotopic mammary fat pad tumors were generated by injecting 5×106 MDA-MB-231 cells (in 1:1 PBS:Matrigel) into the right caudal mammary fat pad of 8-12 week-old, female athymic nu/nu mice (n=30). Mice were obtained from Simonsen Laboratories (Gilroy, CA). Tumors were allowed to grow to ∼100 mm3, at which point they were divided into 6 groups of 5 mice with approximately the same tumor burden and treatment was initiated. Group 1 was vehicle (DMSO) control, group 2 received 5 days of 15 mg/kg paclitaxel, group 3 received 19 days of 60 mg/kg AGX51 bid, group 4 received a combination of paclitaxel and 6.7 mg/kg AGX51 bid, group 5 received a combination of paclitaxel and 20 mg/kg AGX51 bid, group 6 received a combination of paclitaxel and 60 mg/kg bid and group 7 received a combination of paclitaxel and 60 mg/kg AGX51 but only received AGX51 for 7 days (all other AGX51 groups had 19 days of AGX51 treatment). The AGX51 and paclitaxel were administered i.p. Tumor volumes were determined throughout the study using a digital caliper and the formula: tumor volume = ½ (length × width2) where the greatest longitudinal diameter is the length of the tumor and the greatest transverse diameter is the width. At study termination the mice were sacrificed by cervical dislocation.

Lung metastases were generated by injecting 6-8 week-old, female, Balb/c mice with 50,000 luciferase-labeled 4T1 cells into the tail vein. Twenty-four hours after tail vein injections mice were treated with DMSO vehicle or 50 mg/kg AGX51 qd (five mice per treatment group) by i.p. injection. One mouse in the AGX51-treatment group died from the isoflurane anesthesia during the first luciferase imaging session. In another experiment mice were treated bid. The bid experiment was performed three times. In the first experiment n=5 mice per group, in the second experiment n=4 vehicle-treated mice and n=8 AGX51-treated mice and in the third experiment n=5 mice per group. No statistically significant differences in the effect was observed across experiments. Development of lung metastases was monitored using the IVIS-200 *in vivo* imaging system. In the experiment testing the effects of AGX51 on established lung metastases (n=5 mice per group), once evidence of lung metastases was observable by *in vivo* imaging, mice were divided into four groups of five mice with approximately the same tumor burden per treatment group. The groups were: Group 1 vehicle, group 2 50 mg/kg AGX51 bid, group 3 15 mg/kg paclitaxel qd for 5 days, and group 4 combination of AGX51+paclitaxel. At the end of the experiments mice were euthanized and tissues were collected for further analyses. Lung tumor burden was quantified in a blinded fashion by a pathologist.

To assess effects of AGX51 on seeding of the lungs, 20 6-8 week old female Balb/c mice were injected with GFP-labeled 4T1 cells as described above and the next day treated with AGX51 (50 mg/kg) (n=10) or DMSO (n=10). Twenty-four hours later, one set of mice (5 from the AGX51 group and 5 from the DMSO group) was euthanized and their lungs collected for histological analysis. After 48 hours of treatment the remaining 10 mice were euthanized and their lungs collected for histological analysis.

To assess whether *in vivo* treatment results in acquired resistance to AGX51 10 6-8 week old female Balb/c mice were tail vein-injected with GFP-labeled 4T1 cells and treated with AGX51 or DMSO for two weeks as described above. After two weeks of treatment the mice were euthanized and the lungs were dissociated according to the kit instructions (Mouse Lung Dissociation Kit, MACS Miltenyi Biotec) and the cells were sorted for the GFP positive fraction. The cells were allowed to expand to generate enough cells for AGX51 treatment and Western blot analysis, at which point the cells were harvested and whole cell lysates were made.

Spontaneous colon tumors were induced by treating 30, 4-week old male A/J mice (Jackson Laboratory) with AOM (10 mg/kg; Sigma Aldrich) once a week by i.p. injection for 6 weeks. Mice were maintained on AIN-93G purified diet (Research Diets) for the duration of the experiment. After a three-week treatment break mice were treated i.p. with DMSO (n=13) or AGX51 (n=11) (15 mg/kg) bid for three weeks. Following the last injection, the mice were euthanized and colon tumors were formalin fixed to assess tumor burden. Tumor numbers and size were determined in whole mounts of the tissues following methylene blue staining.

### Immunohistochemistry

Slides mounted with sections from formalin-fixed and paraffin-embedded mouse tissues were deparaffinized, rehydrated and stained with hematoxylin and eosin (H&E), as well as with the following primary antibodies: Id1 (37-2, Biocheck 1:1000), Vimentin (5741, Cell Signaling 1:500), Cyclin D1 (2978, Cell Signaling 1:500). Detection was performed using the Vectastain Elite ABC kit (Vector), diaminobenzidine was used as the chromogen, and Harris Modified Hematoxylin was used as the nuclear counterstain.

### Immunofluorescence

The lungs from mice from the seeding experiment were perfused with 1X PBS, followed by 4% PFA. The lungs were then removed, washed in cold PBS and fixed in 4% PFA overnight. The lungs were dehydrated in a sucrose solution series (20-30%). The lungs were then embedded in OCT and flash frozen. Ten micron cryosections were then boiled for 10 minutes in citrate buffer, for antigen retrieval, washed with 1X PBS, blocked in PBS, 10% goat serum, 0.05% triton and then incubated with anti-GFP antibody (GFP-1020, Aves labs, Inc.) overnight at 4 degrees. The next day the slides were washed, and incubated with Alexa Fluor 555 anti-chicken antibody (A-21437, Thermo Fisher Scientific), washed and counterstained with DAPI followed by mounting and coverslip application. The slides were scanned and ten random fields from each section were analyzed for GFP positive cell content.

### AGX51 resistant clone generation

To generate resistant cell lines, cells were maintained in AGX51-containing media (20-40 μM range). AGX51-containing media was replaced every 2-3 days. Any clones that grew out were expanded, and maintained in AGX51-containing media. Transiently resistant clones expressing Id1/Id3 had their cDNA sequenced for the coding regions of Id1 and Id3 by Sanger sequencing using the primers:

Id1-F:CTTCTTGTTCTCTTCCCACA
Id1-R:GATCAAACCCTCTACCCACT
Id3-F:CACTGTTTGCTGCTTTAGGT
Id3-R:CGTTGAGTTCAGGGTAAGTG

### Statistical Analyses

Three replicates were generally used for each experimental condition for *in vitro* experiments and 5 mice per group were typically used in each mouse experiment. The sample sizes were determined based on an expected large effect size. With 3 replicated per condition, an effect size as small as 3 can be detected with 80% power at a two-sided significance level of 0.05 using a two-sample t-test. With 5 mice per group, an effect size as small as 2 can be detected with 80% power at a two-sided significance level of 0.05 using a two-sample t-test. Additional experiments may be performed when larger variation in data was observed and data were pooled for analysis. In general, Welch’s t-test was used to examine differences between two groups. ANOVA was used to examine differences across multiple experimental groups. Data may be transformed to ensure the underlying normality assumptions were met. For example, for tumor number data, square root transformation was used. Weighted linear regression analysis was used when heteroscedasticity was observed and data points in each group were typically weighted by the reciprocal of the standard deviation of data in each group. For data pooled from multiple experiments, the model included both experiments and experiments by treatment group interaction as covariates to account for potential differences in experiments. Significance of linear contrasts of interest was assessed based on estimates obtained from the weighted least squares. Q-Q plot of the residuals was examined to ensure the underlying model assumptions were met. P-value <0.05 was considered statistically significant. For the colon tumor counts, the data was analyzed using the Wilcoxon Rank Sum test.

## Supplemental Information Titles and Legends

**Figure S1.**
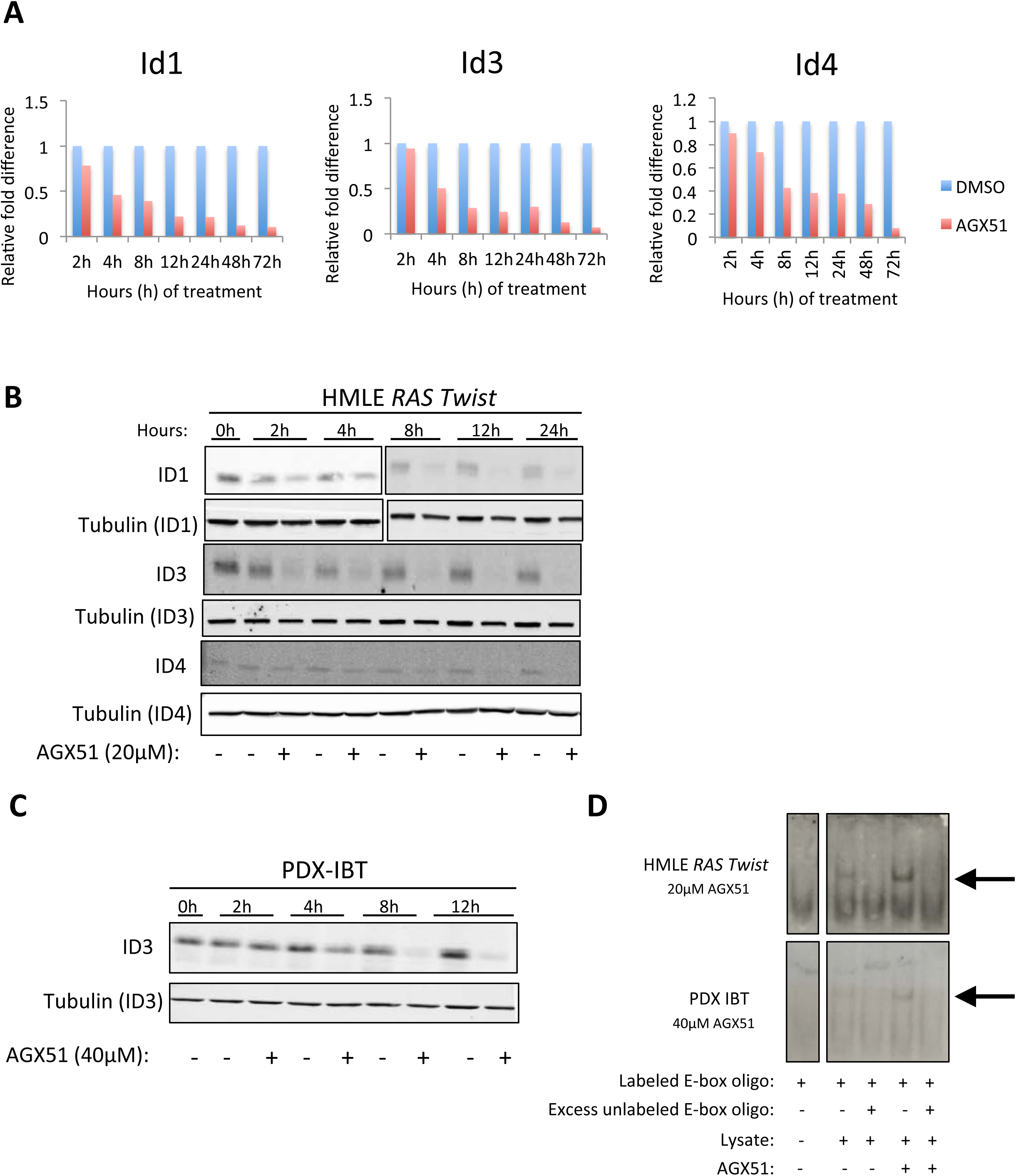
Protein expression and EMSAs in HMLE *RAS Twist* and PDX IBT cell lysates. Related to Figure 1. (A) Quantification of Western blot data presented in Figure 1b. (B) Western blot for Id1, Id3 and Id4 on whole cell lysates from HMLE *RAS Twist* cells treated with 20 µM AGX51 for 0-24 hours. (C) Western blot for Id3 on whole cell lysates from PDX IBT cells treated with 40 µM AGX51 for 0-12 hours. (D) EMSA on lysates from HMLE *RAS Twist* (upper blots) and PDX IBT (lower blots) cells treated with 20 or 40 µM AGX51, respectively, for 24 hours. Arrows indicate binding to DNA. Tubulin is used as a protein loading control.

**Figure S2.**
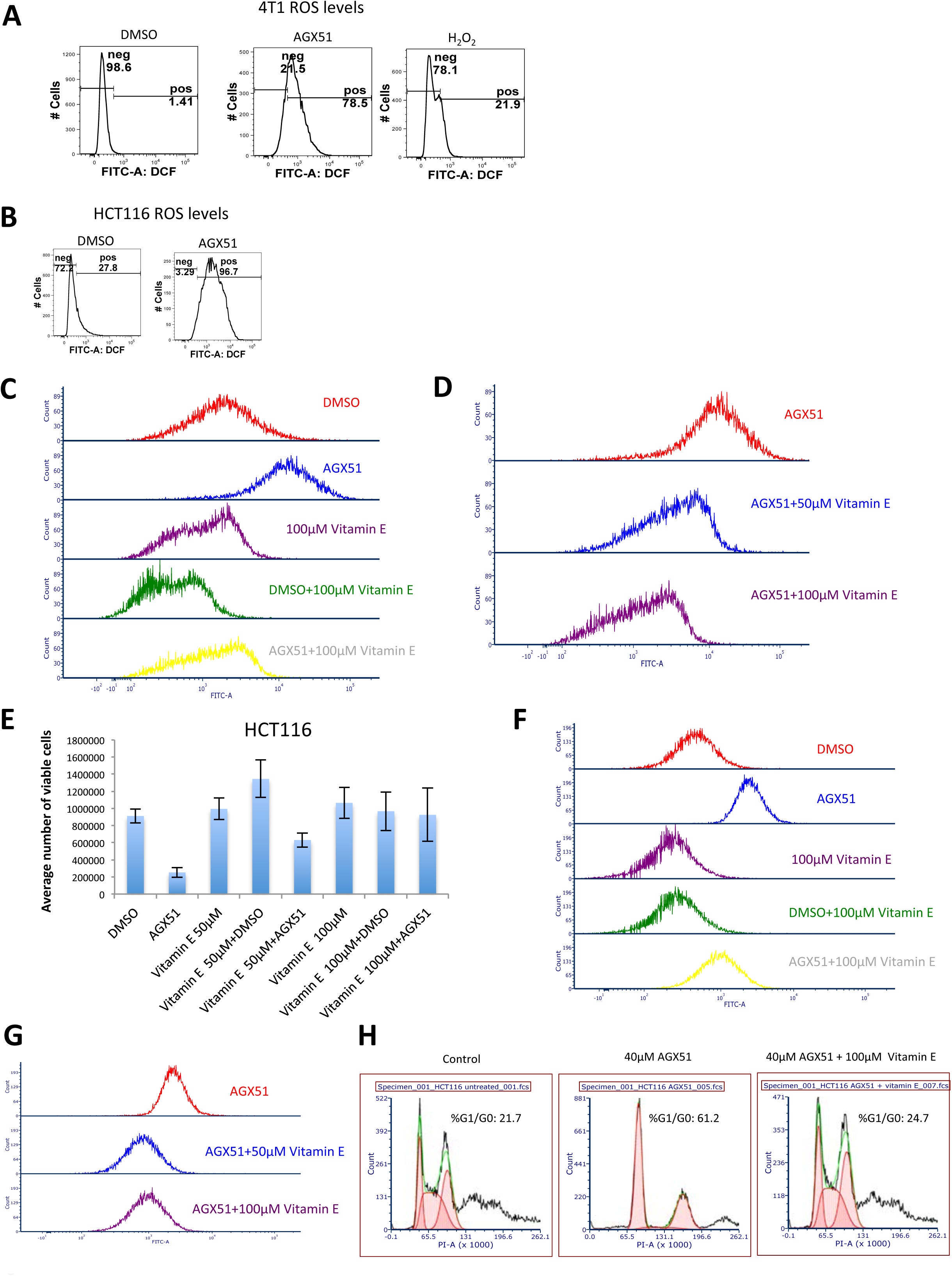
ROS levels in HCT116 and 4T1 cells following AGX51 treatment with and without vitamin E. Related to Figure 2. (A) Reactive oxygen species accumulation in 4T1 cells induced by 24 hour treatment with AGX51 (40 µM) or H_2_O_2_ (100 µM), as determined by H2DCFDA staining followed by flow cytometry analysis, with FITC positivity indicative of ROS presence. (B) Reactive oxygen species accumulation in HCT116 cells induced by AGX51 treatment (C) Reactive oxygen species levels in 4T1 cells treated with 40 µM AGX51 for 24 hours with or without 1 hour vitamin E pre-treatment (100 µM). (D) Reactive oxygen species levels in 4T1 cells treated with 40 µM AGX51 for 24 hours with or without 1 hour vitamin E pre-treatment (50 or 100 µM). (E) Viable cell numbers, as determined by trypan blue exclusion and cell counting, of HCT116 cells pretreated with vitamin E (50 or 100 µM) for 1 hour followed by 24 hours of AGX51 (40 µM), or DMSO control. (F) Reactive oxygen species levels in HCT116 cells treated with 40 µM AGX51 for 24 hours with or without 1 hour vitamin E pre-treatment (100 µM). (G) Reactive oxygen species levels in HCT116 cells treated with 40 µM AGX51 for 24 hours with or without 1 hour vitamin E pre-treatment (50 or 100 µM). (H) Cell cycle analysis of HCT116 cells treated with 40 µM AGX51 for 24 hours, with or without 1 hour pretreatment with 100 µM vitamin E.

**Figure S3.**
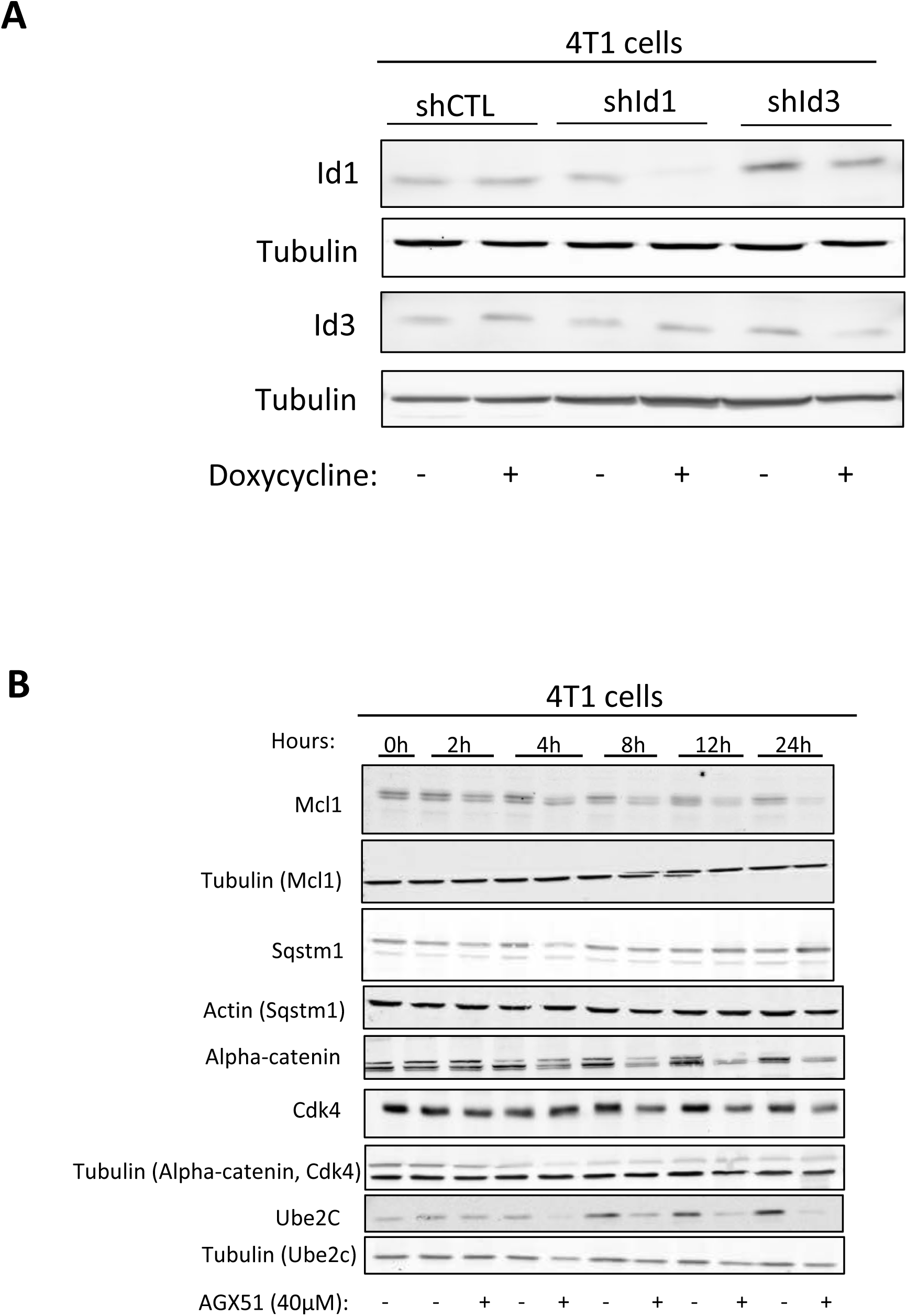
Protein expression in 4T1 cells. Related to Figure 4. (A) Western blot for Id1 and Id3 on whole cell lysates from 4T1 cells transduced with doxycycline-inducible constructs expression short hairpins against Renilla (shCTL), Id1 (shId1) or Id3 (shId3) with or without doxycycline treatment. (B) Western blot for Mcl1, Sqstm1, Alpha-catenin, Cdk4, and Ube2C on whole cell lysates from 4T1 cells treated with 40 µM AGX51 for 0-24 hours.

**Figure S4.**
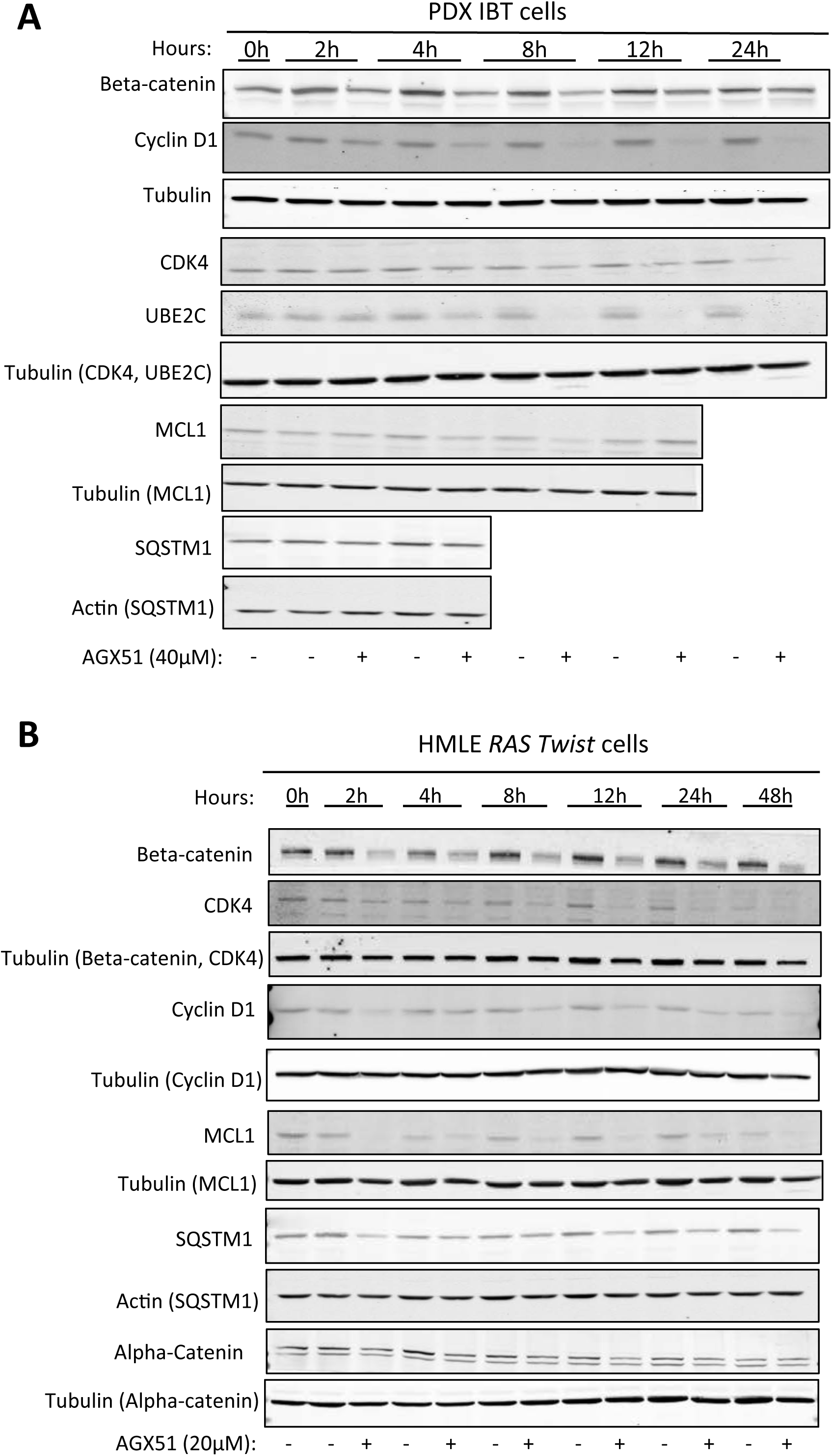
Protein expression in PDX IBT and HMLE *RAS Twist* cells. Related to Figure 4. (A) Western blot for Beta-catenin, Cyclin D1, CDK4, UBE2C, MCL1 and SQSTM1 on whole cell lysates from PDX IBT cells treated with 40 µM AGX51 for 0-24 hours. Tubulin/Actin are used as protein loading controls. (B) Western blot for Beta-catenin, Cyclin D1, CDK4, SQSTM1, MCL1 and Alpha-catenin on whole cell lysates from HMLE *RAS Twist* cells treated with 20 µM AGX51 for 0-48 hours.

**Figure S5.**
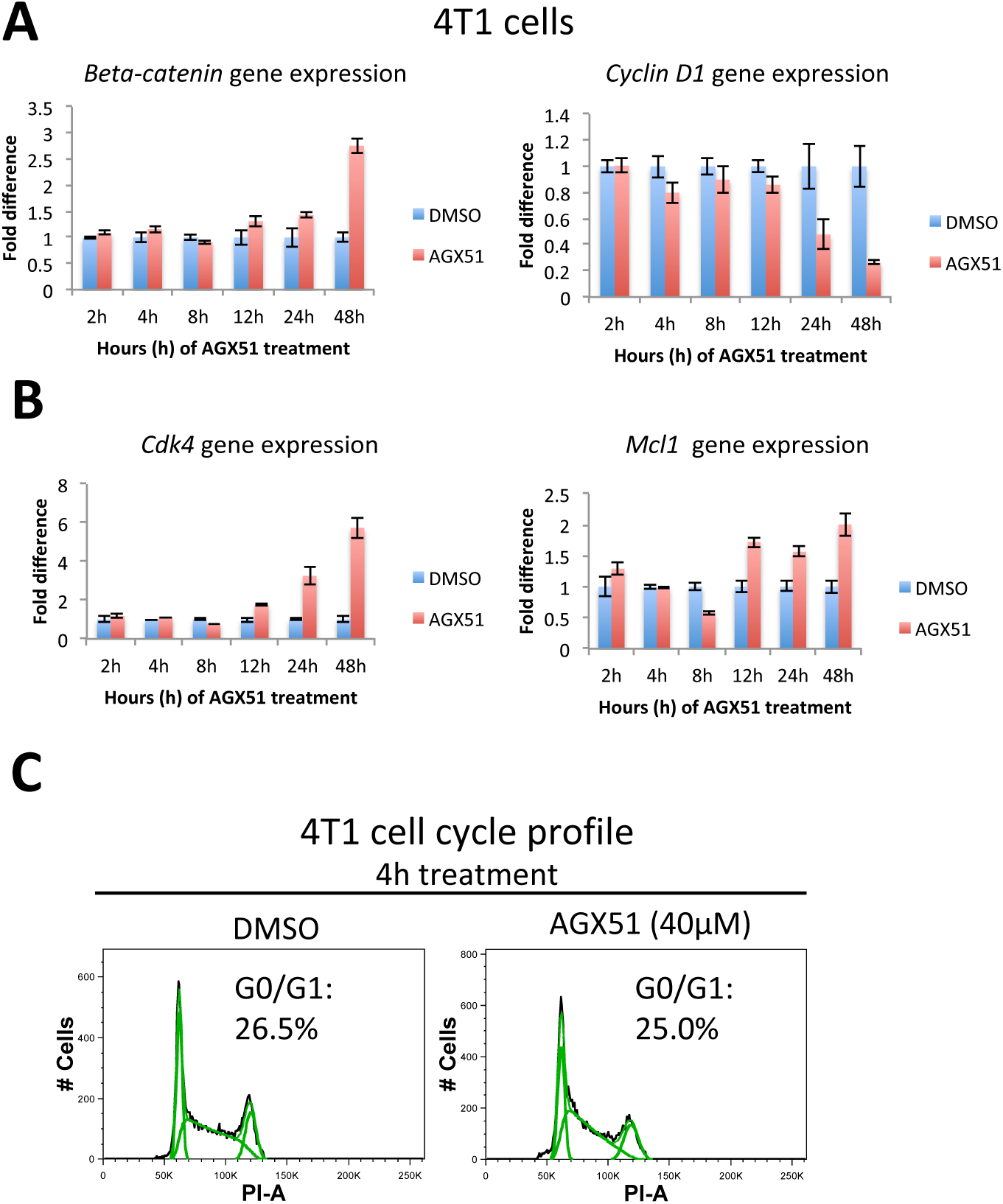
Gene expression analysis of genes found differentially expressed in SILAC analysis of AGX51-treated 4T1 cells. Related to Figure 4. (A) qRT-PCR analysis for relative *Beta-catenin* and *Cyclin D1 mRNA levels* in 4T1 cells treated with 40 µM AGX51 for 2-48 hours. Data shows mean of technical triplicates with error bars representing SEM. (B) qRT-PCR analysis for *Cdk4* and *Mcl1* expression in 4T1 cells treated with 40 µM AGX51 for 2-48 hours. Data shows mean of technical triplicates with error bars representing SEM. (C) Cell cycle analysis of 4T1 cells treated with 40 µM AGX51 for four hours.

**Figure S6.**
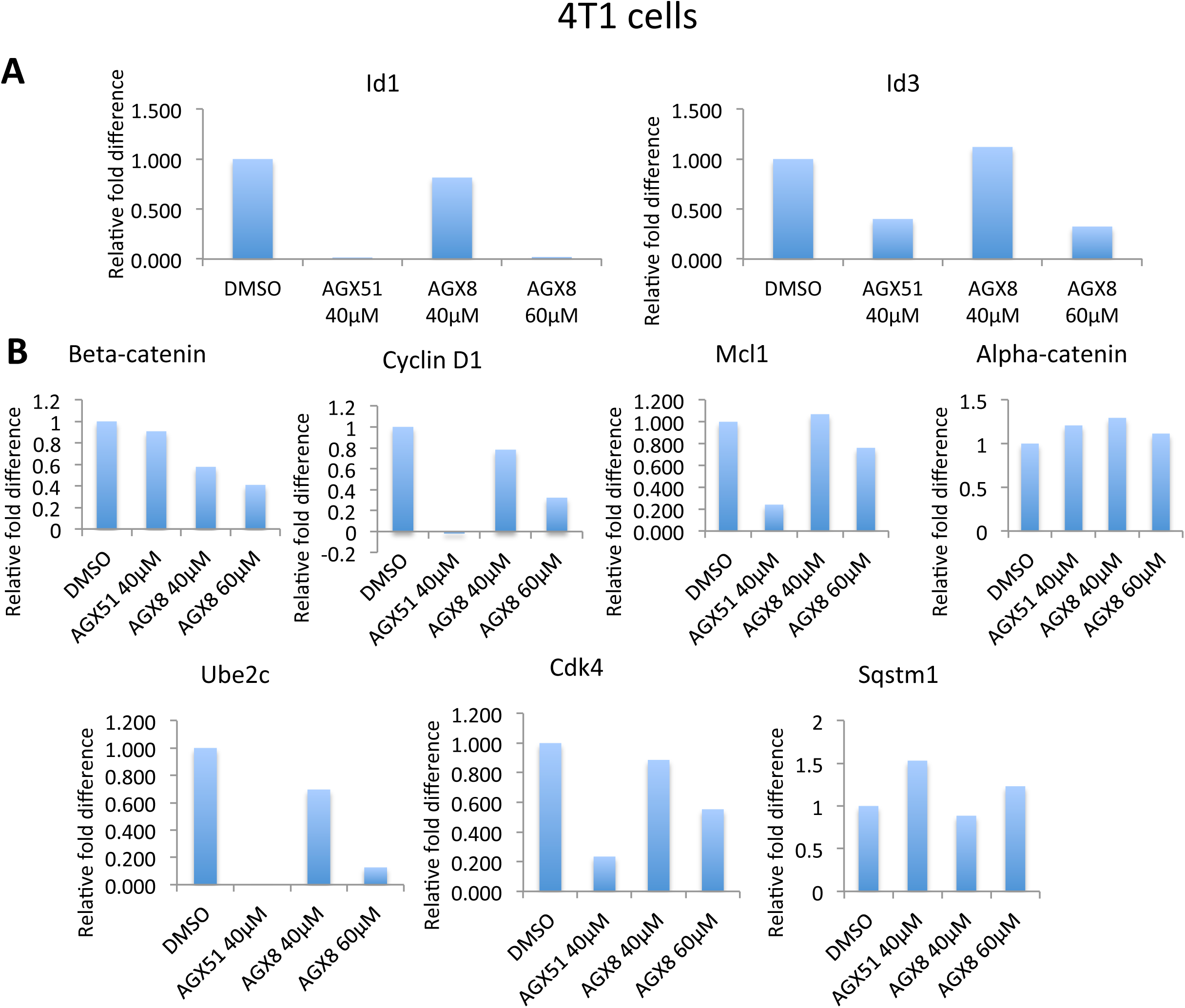
Western blot quantification. Related to Figure 4. (A and B) Western blot quantification for Western blots presented in Figure 4G and H, respectively.

**Figure S7.**
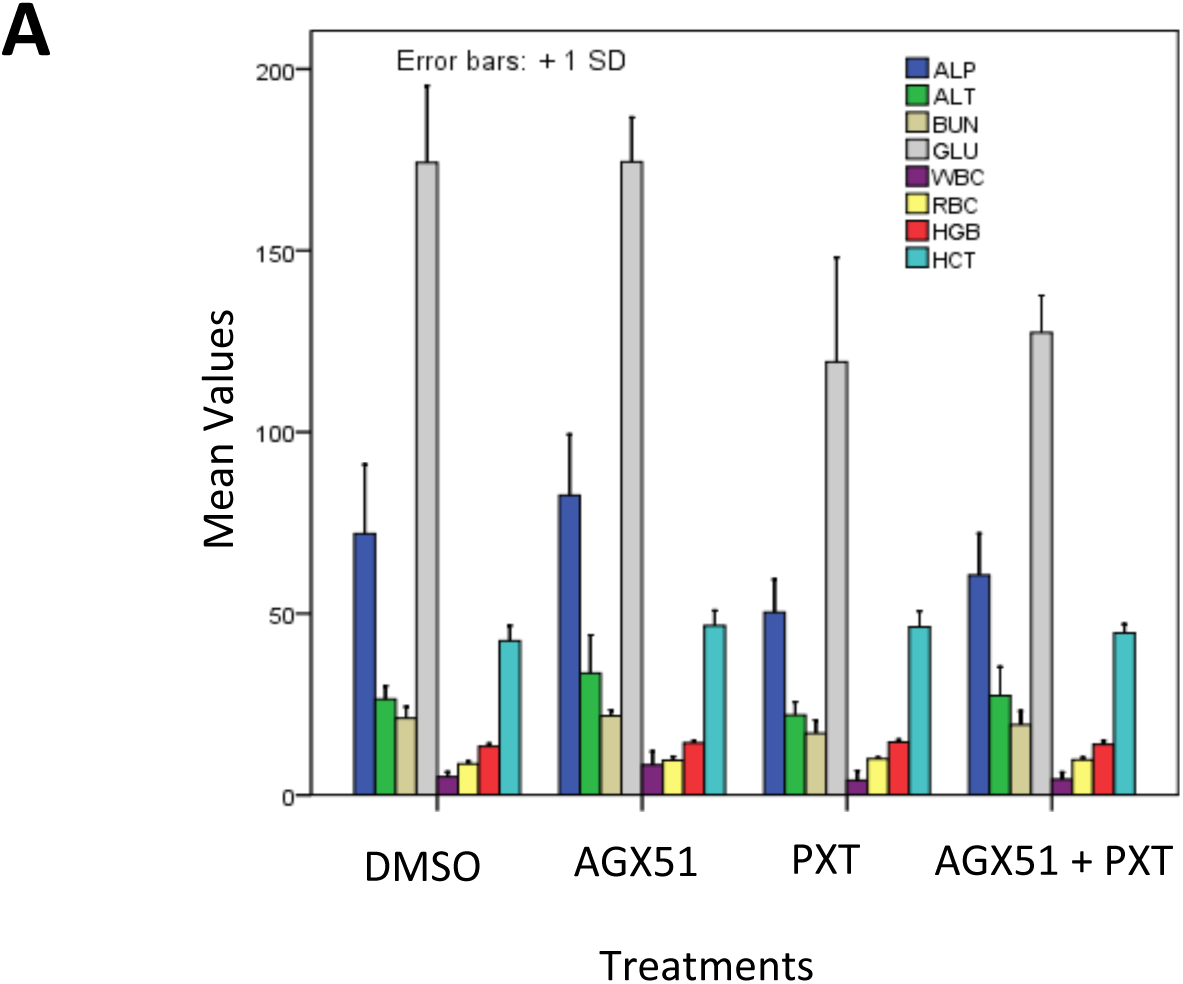
*In vivo* toxicity analysis of AGX51. Related to Figure 5. (A) Systemic clinical chemistry and hematology concentrations were determined in mice treated for 14 days with vehicle, AGX51 (60 mg/kg, qd), paclitaxel (15 mg/kg for 5 days), or paclitaxel plus AGX51 (n=5 mice per group). Mean values (with error bars showing SD) for ALP, alkaline phosphatase; ALT, alanine aminotransferase; BUN, blood urea nitrogen; GLU, glucose; WBC, white blood cell count; RBC, red blood cell count; HGB, hemoglobin; HCT, hematocrit are shown. Compared to vehicle, no effect of the AGX51 treatment on clinical chemistry or hematology values was observed. Similarly, values from the paclitaxel group and the AGX51+paclitaxel were statistically similar.

**Figure S8.**
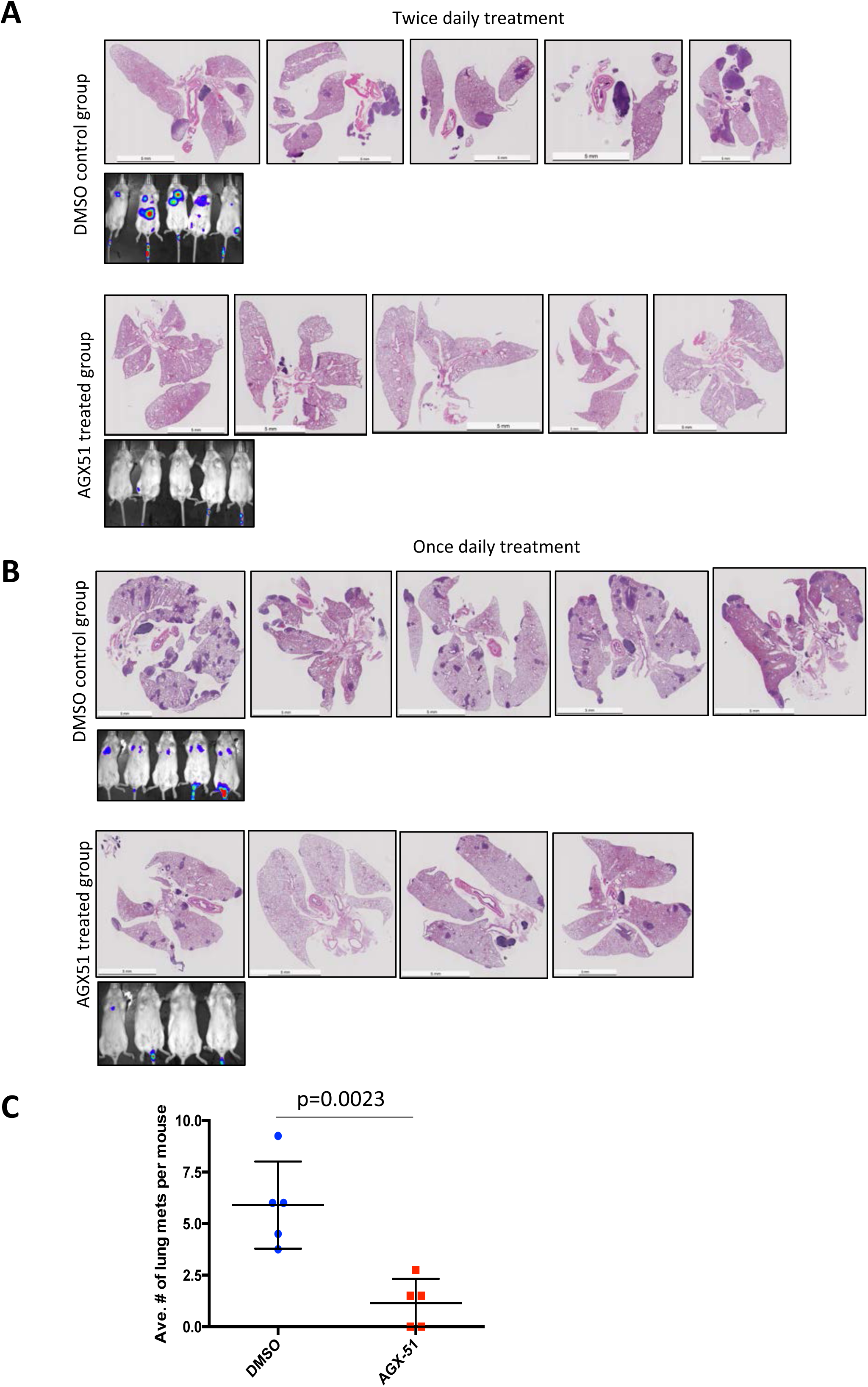
Lung metastases in AGX51 treated mice. Related to Figure 5. (A) H&E staining, and corresponding bioluminescent imaging from mice injected with 5×104 4T1 cells via tail vein and treated with 50 mg/kg bid or vehicle (n=5 mice per treatment group). (B) H&E staining, and corresponding bioluminescent imaging from mice injected with 5×106 4T1 cells via tail vein and treated with 50 mg/kg qd or vehicle (n=5 mice per treatment group, where one mouse died in the AGX51 treatment group following isoflurane anesthesia during the first bioluminescent imaging session). (C) Average number of lung metastases in mice injected with 5×104 4T1 cells via tail vein, treated with 50 mg/kg AGX51 qd or vehicle (n=5 or n=5, respectively). Means are plotted with error bars showing SEM. AGX51 treatment had a significant effect on lung tumor development (p<0.0023).

**Figure S9.**
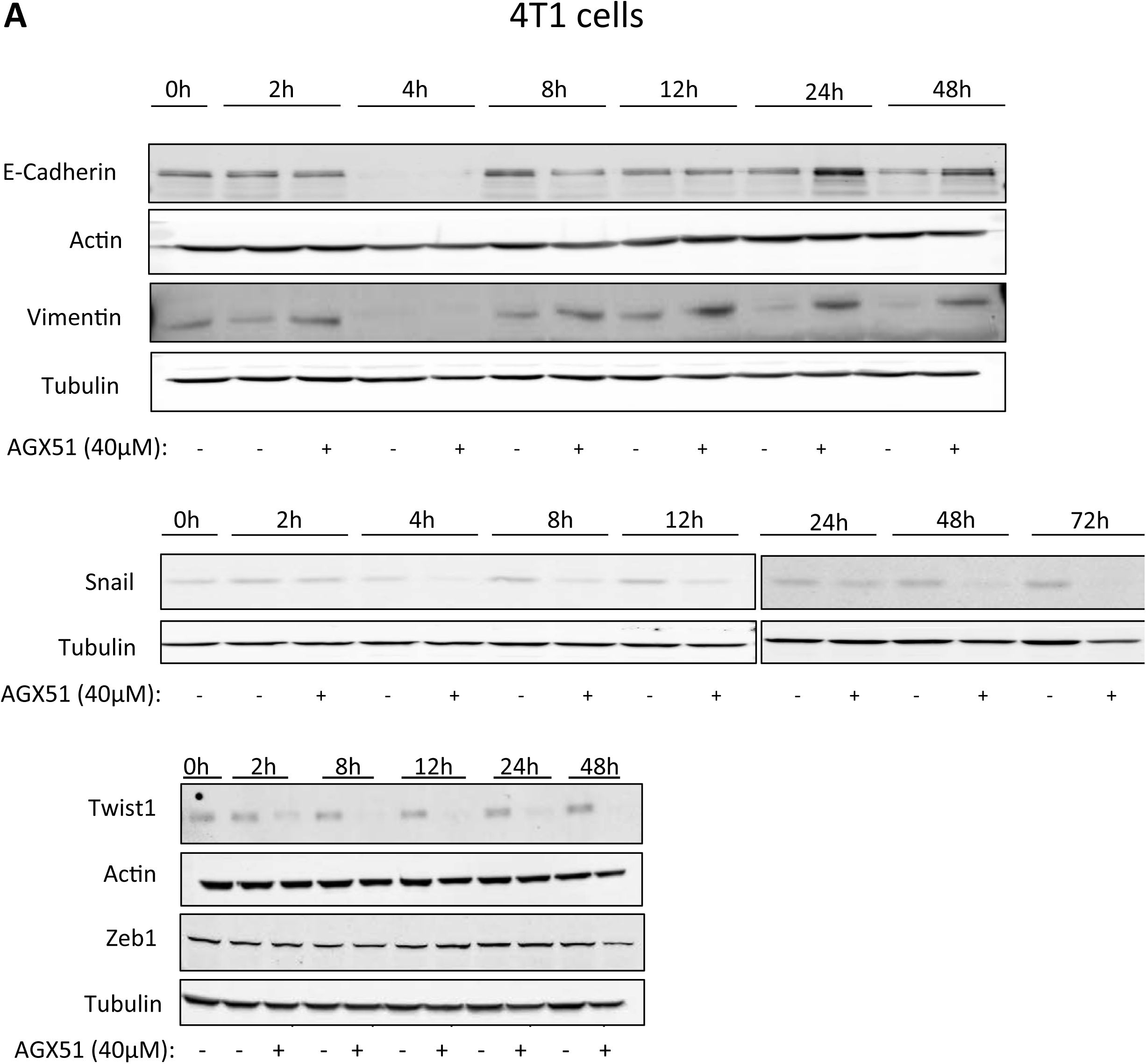
Effects of AGX51 on EMT signatures. Related to Figure 4. (A) Western blot for E-cadherin, Vimentin, Snail, Twist1 and Zeb1 on whole cell lysates from 4T1 cells treated with 40 µM AGX51 for 0-48, or 0-72 hours. Tubulin/Actin serve as protein loading controls.

**Figure S10.**
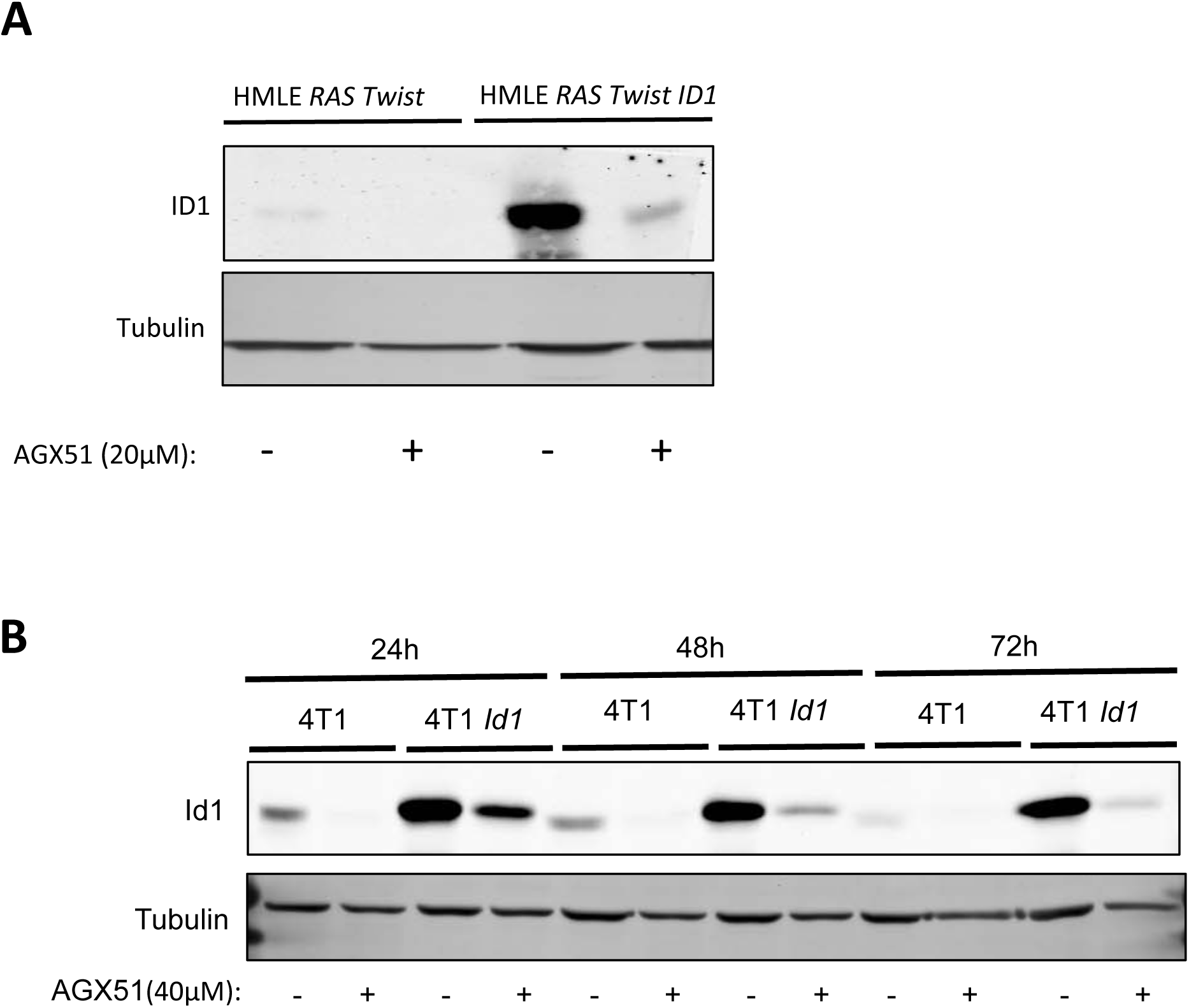
Effects of AGX51 on Id1-overexpressing cells. Related to Figure 6. (A) Western blot for Id1 on whole cell lysates from HMLE *RAS Twist* and HMLE *RAS Twist ID1* cells treated with 20 µM AGX51 for 24 hours. (B) Western blot for Id1 on whole cell lysates from 4T1 and 4T1 *Id1* cells treated with 40 µM AGX51 for 24, 48 and 72 hours.

**Figure S11.**
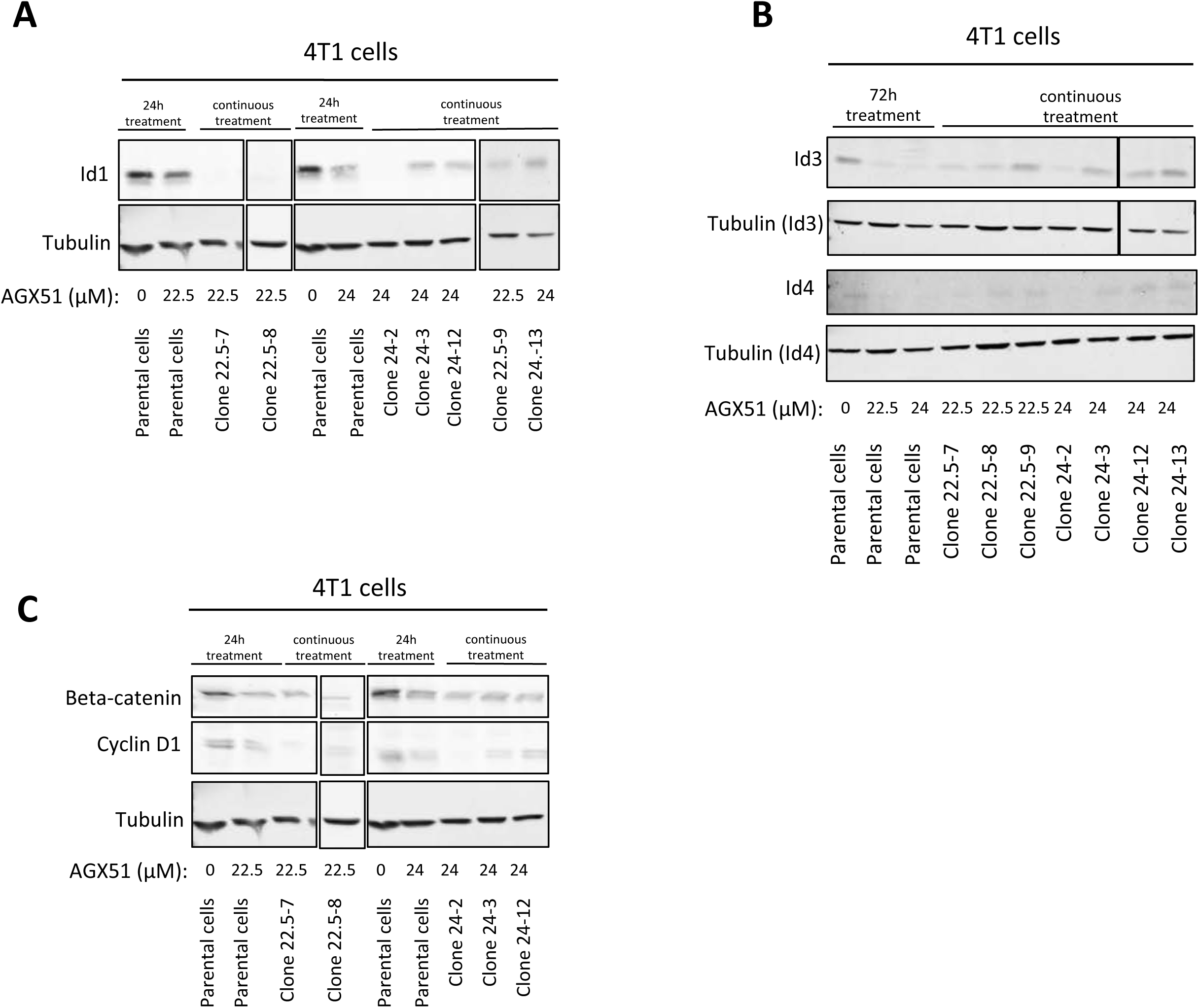
Beta-catenin and Cyclin D1 expression in AGX51 transiently resistant clones. Related to Figure 6. (A) Western blot for Id1 on parental 4T1cells (treated for 24 hours) and transiently resistant clones treated (continuously) with 22.5 or 24 µM AGX51. (B) Western blot for Id3 and Id4 on parental 4T1 cells (treated for 72 hours) and transiently resistant clones treated (continuously) with 22.5 or 24 µM AGX51. (C) Western blot analysis for Beta-catenin and Cyclin D1 on cell lysates from parental 4T1 cells or transiently resistant clones treated with 22.5 or 24 µM AGX51 for 24 hours or continuously, as indicated.

---

**Table S1.** IC50 values, as determined by MTT assay, for AGX51-treated cell lines. Related to Figure 1.

| Cell Type | Cell line | IC50 value after 24 hours of treatment |
| --- | --- | --- |
| cell line | 4T1 | 26.66 μM |
| cell line | HMLE *RAS Twist* | 8.7 μM |
| TNBC cell line | MDA-MB-157 | 22.28 μM |
| | MDA-MB-436 | 30.91 μM |
| | MDA-MB-231 | NA (60 μM=70%) |
| HER-2 +ve cell line | MDA-MB-453 | NA (60 μM=57%) |
| | BT-474 | NA (60 μM=53%) |
| | MDA-MB-361 | NA (60 μM=53%) |
| | SK-BR-3 | 36.55 μM |
| ER +ve cell line | MCF-7 | 60 μM |
| | T47-D | NA (60 μM=65%) |
| PDX cell line | PDX-BR7 | 10.89 μM |
| | PDX-IBT | 11.97 μM |
| | PDX-BR11 | 18.56 μM |
| cell line | 4T1 i.shRNA-*Id1* (-dox.) | 25.40 μM |
| cell line | 4T1 i.shRNA-*Id1* (+dox.) | 9.03 μM |
| cell line | 4T1 i.shRNA-*Id3* (-dox.) | 24.98 μM |
| cell line | 4T1 i.shRNA-*Id3* (+dox.) | 8.55 μM |

Legend: +, positive; TNBD, Triple negative breast cancer; ER, estrogen receptor; PDX, patient derived xenograft; i.shRNA, inducible shRNA; dox., doxycyclin

**Table S2.** Proteins differently expressed by SILAC in 4T1 cells treated with AGX51 for 4 hours. Related to Figure 3.

| Gene | Protein | Molecular Weight | Fold difference |  |  |
| --- | --- | --- | --- | --- | --- |
|  |  |  | DMSO (Heavy) vs. AGX51 | DMOS (Light1) vs. AGX (Heavy1) | DMOS (Light2) vs. AGX (Heavy2) |
| *Zbed5* | B2RPU8 | 84 kDa | -2.78 | -2.85 | -2.64 |
| *Sqstm1* | SQSTM1 | 44 kDa | -2.36 | -2.43 | -3.17 |
| *Ctnnb1* | CTNNB1 | 85 kDa | -2.27 | -1.35 | -1.41 |
| *Morf4l2* | MO4L2 | 32 kDa | -2.24 | -1.57 | -1.55 |
| *Ccnd1* | CCND1 | 33 kDa | -2.17 | -1.62 | -3.1 |
| *Mcl1* | MCL1 | 35 kDa | -2.14 | -1.77 | -2.01 |
| *Ctnna1* | CTNA1 | 100 kDa | -1.63 | -2.03 | -1.94 |
| *Xirp2* | XIRP2 | 428 kDa | -1.63 | -1.97 | -2.3 |
| *Paf* | PAF15 | 12 kDa | -1.51 | -1.79 | -1.71 |
| *Asf1b* | ASF1B | 22 kDa | -1.5 | -1.39 | -1.55 |
| *Cdk4* | CDK4 | 34 kDa | -1.48 | -1.35 | -1.44 |
| *Uck2* | UCK2 | 29 kDa | -1.44 | -1.35 | -1.63 |
| *Id1* | A2AHY3 | 18 kDa | -1.39 | -2.07 | -1.71 |
| *Ube2c* | UBE2C | 20 kDa | -1.39 | -1.5 | -1.59 |
| *Vim* | VIME | 54 kDa | 1.36 | 3.52 | 1.36 |
| *Lonp1* | LONM | 106 kDa | 1.38 | 2.24 | 1.96 |
| *Phb* | PHB | 30 kDa | 1.38 | 4.21 | 1.97 |
| *Pth2* | PTH2 | 20 kDa | 1.38 | 3.17 | 1.56 |
| *Pam16* | TIM16 | 14 kDa | 1.39 | 2.8 | 1.52 |
| *Suc1a2* | SUCB1 | 50 kDa | 1.39 | 2.01 | 1.48 |
| *Cox4i1* | COX41 | 20 kDa | 1.44 | 3.02 | 1.67 |
| *Gnl3* | GNL3 | 61 kDa | 1.44 | 1.5 | 1.35 |
| *Acaa2* | THIM | 42 kDa | 1.5 | 2.02 | 1.67 |
| *Hadha* | ECHA | 83 kDa | 1.5 | 2.39 | 1.68 |
| *Ak4* | KAD4 | 25 kDa | 1.51 | 1.46 | 1.35 |
| *Vdac1* | VDAC1 | 31 kDa | 1.53 | 4.29 | 1.67 |
| *Acot13* | ACO13 | 15 kDa | 1.54 | 2.25 | 1.62 |
| *Mrps6* | RT06 | 14 kDa | 1.55 | 1.89 | 1.41 |
| *Fxn* | FRDA | 23 kDa | 1.55 | 1.69 | 1.47 |
| *Aco2* | ACON | 85 kDa | 1.56 | 1.98 | 1.59 |
| *Pdhb* | ODPB | 39 kDa | 1.58 | 1.64 | 1.6 |
| *Hspa9* | GRP75 | 73 kDa | 1.58 | 2.12 | 1.91 |
| *Lrpprc* | LPPRC | 157 kDa | 1.59 | 2.39 | 1.5 |
| *Idh3a* | IDH3A | 40 kDa | 1.61 | 2.06 | 1.63 |
| *Gcat* | KBL | 45 kDa | 1.62 | 2.1 | 1.49 |
| *Pck2* | PCKGM | 71 kDa | 1.62 | 2.14 | 1.67 |
| *Slirp* | SLIRP | 11 kDa | 1.62 | 2.68 | 1.66 |
| *Mdh2* | MDHM | 36 kDa | 1.63 | 2.12 | 1.78 |
| *Got2* | AATM | 47 kDa | 1.63 | 2.1 | 1.66 |
| *Grpel1* | GRPE1 | 24 kDa | 1.63 | 1.96 | 1.48 |
| *Acat1* | THIL | 45 kDa | 1.63 | 1.98 | 1.48 |
| *Sod2* | SODM | 25 kDa | 1.66 | 2.11 | 1.62 |
| *Eci1* | ECI1 | 32 kDa | 1.67 | 2.36 | 1.54 |
| *Cs* | CISY | 52 kDa | 1.67 | 2.02 | 1.45 |
| *Sdha* | SDHA | 73 kDa | 1.68 | 2.07 | 1.52 |
| *Idh2* | IDHP | 51 kDa | 1.68 | 2.22 | 1.56 |
| *Aldh2* | ALDH2 | 57 kDa | 1.69 | 2.49 | 1.73 |
| *Acadl* | ACADL | 48 kDa | 1.71 | 2.7 | 1.83 |
| *Hadhb* | ECHB | 51 kDa | 1.72 | 2.73 | 1.49 |
| *Shmt2* | Q9CZ7 | 56 kDa | 1.74 | 2.3 | 1.66 |
| *Trap1* | TRAP1 | 80 kDa | 1.74 | 1.91 | 1.42 |
| *Mthfd1l* | C1TM | 106 kDa | 1.75 | 2.21 | 1.65 |
| *Hspd1* | CH60 | 61 kDa | 1.75 | 2.42 | 1.98 |
| *Aldh18a1* | P5CS | 87 kDa | 1.75 | 1.98 | 2.17 |
| *Atp5d* | ATPD | 18 kDa | 1.75 | 2.8 | 1.78 |
| *Mrps34* | RT34 | 26 kDa | 1.75 | 2.67 | 1.78 |
| *Idh3b* | Q91VA7 | 42 kDa | 1.76 | 2.73 | 1.64 |
| *Atp5b* | ATPB | 56 kDa | 1.78 | 2.95 | 1.97 |
| *Idh3g* | IDHG1 | 43 kDa | 1.78 | 1.86 | 1.72 |
| *Atp5a1* | ATPA | 60 kDa | 1.79 | 3.41 | 1.95 |
| *Acot9* | ACOT9 | 51 kDa | 1.79 | 2.2 | 1.55 |
| *Mcoln2* | MCLN2 | 65 kDa | 1.79 | 1.49 | 2.01 |
| *110037F02R1* | E9PZY8 | 207 kDa | 1.8 | 1.8 | 1.56 |
| *C1qbp* | Q8R5L1 | 31 kDa | 1.83 | 2.47 | 1.79 |
| *Mrps23* | RT23 | 20 kDa | 1.83 | 1.38 | 1.74 |
| *Hsd17b10* | Q99N15 | 27 kDa | 1.83 | 2.3 | 1.64 |
| *Hspe1* | CH10 | 11 kDa | 1.84 | 2.14 | 1.78 |
| *Dld* | DLDH | 54 kDa | 1.85 | 2.26 | 1.62 |
| *Me2* | MAOM | 66 kDa | 1.85 | 3.14 | 1.86 |
| *Atp5j2* | ATPK | 10 kDa | 1.92 | 5.64 | 2.23 |
| *Mrps16* | RT16 | 15 kDa | 1.99 | 2.5 | 1.78 |
| *Mrps17* | RT17 | 13 kDa | 2.02 | 2.33 | 1.93 |
| *Mrpl12* | RM12 | 22 kDa | 2.03 | 2.33 | 1.69 |
| *Mrpl40* | D3Z7C0 | 19 kDa | 2.04 | 2.13 | 1.69 |
| *Prdx3* | PRDX3 | 28 kDa | 2.06 | 2.39 | 1.89 |
| *Dlat* | ODP2 | 68 kDa | 2.1 | 2.55 | 1.96 |
| *Mrpl53* | RM53 | 13 kDa | 2.12 | 2.43 | 1.81 |
| *Dlst* | ODO2 | 49 kDa | 2.13 | 2.01 | 2.04 |
| *Cox5a* | COX5A | 16 kDa | 2.15 | 2.75 | 2.02 |
| *Glud1* | DHE3 | 61 kDa | 2.17 | 2.01 | 1.88 |
| *Atp5o* | ATPO | 23 kDa | 2.25 | 4.4 | 1.96 |
| *Atp5h* | ATP5H | 19 kDa | 2.36 | 5.02 | 2.45 |
| *Nbas* | E9Q411 | 266 kDa | 7.01 | 2.58 | 1.56 |
| *Ifrd1* | IFRD1 | 50 kDa | 1536.69 | 1.95 | 2.34 |

**Table S3.** IC50 values, as determined by MTT assay, for AGX51- and AGX8-treated cell lines. Related to Figure 3.

| Cell line | IC50 value (24 hours) |  |
| --- | --- | --- |
|  | AGX51 | AGX8 |
| 4T1 | 26.66 μM | 32.68 μM |
| HMLE *RAS Twist* | 8.7 μM | 42.25 μM |
| 4T1 i.shRNA-*Id1* (-dox.) | 25.40 μM | 39.49 μM |
| 4T1 i.shRNA-*Id1* (+dox.) | 9.03 μM | 38.77 μM |
| 4T1 i.shRNA-*Id3* (-dox.) | 24.98 μM | 42.19 μM |
| 4T1 i.shRNA-*Id3* (+dox.) | 8.55 μM | 41.59 μM |

Legend: i.shRNA, inducible shRNA; dox., doxycyclin
Note: AGX51 data is also shown in Extended Data Table 1

**Table S4.** Final tumor volumes of MDA-MB-231 tumors treated with paclitaxel, AGX51, paclitaxel + AGX51 or DMSO.

| **Treatment group** | **Tumor volume (mm3)** |
| --- | --- |
| DMSO | 683.91 |
| DMSO | 492.82 |
| DMSO | 411.77 |
| DMSO | 400.60 |
| DMSO | 413.37 |
| 60mg/kg AGX51 (19 days) | 527.73 |
| 60mg/kg AGX51 (19 days) | 647.84 |
| 60mg/kg AGX51 (19 days) | 410.90 |
| 60mg/kg AGX51 (19 days) | 307.18 |
| 60mg/kg AGX51 (19 days) | 373.24 |
| 15mg/kg Paclitaxel+60mg/kg AGX51 (19 days) | 161.33 |
| 15mg/kg Paclitaxel+60mg/kg AGX51 (19 days) | 70.59 |
| 15mg/kg Paclitaxel+60mg/kg AGX51 (19 days) | 88.88 |
| 15mg/kg Paclitaxel+60mg/kg AGX51 (19 days) | 50.34 |
| 15mg/kg Paclitaxel+60mg/kg AGX51 (19 days) | 59.00 |
| 15mg/kg Paclitaxel+20mg/kg AGX51 (19 days) | 182.17 |
| 15mg/kg Paclitaxel+20mg/kg AGX51 (19 days) | 295.06 |
| 15mg/kg Paclitaxel+20mg/kg AGX51 (19 days) | 79.96 |
| 15mg/kg Paclitaxel+20mg/kg AGX51 (19 days) | 143.02 |
| 15mg/kg Paclitaxel+20mg/kg AGX51 (19 days) | 42.15 |
| 15mg/kg Paclitaxel+6.7mg/kg AGX51 (19 days) | 206.13 |
| 15mg/kg Paclitaxel+6.7mg/kg AGX51 (19 days) | 165.44 |
| 15mg/kg Paclitaxel+6.7mg/kg AGX51 (19 days) | 184.99 |
| 15mg/kg Paclitaxel+6.7mg/kg AGX51 (19 days) | 142.83 |
| 15mg/kg Paclitaxel+6.7mg/kg AGX51 (19 days) | 25.81 |
| 15mg/kg Paclitaxel+60mg/kg AGX51 (7 days) | 142.16 |
| 15mg/kg Paclitaxel+60mg/kg AGX51 (7 days) | 109.29 |
| 15mg/kg Paclitaxel+60mg/kg AGX51 (7 days) | 96.58 |
| 15mg/kg Paclitaxel+60mg/kg AGX51 (7 days) | 124.37 |
| 15mg/kg Paclitaxel+60mg/kg AGX51 (7 days) | 50.97 |
| 15mg/kg Paclitaxel | 305.21 |
| 15mg/kg Paclitaxel | 152.97 |
| 15mg/kg Paclitaxel | 267.26 |
| 15mg/kg Paclitaxel | 259.79 |

## References

Alcantara Llaguno, S., Chen, J., Kwon, C.H., Jackson, E.L., Li, Y., Burns, D.K., Alvarez-Buylla, A., and Parada, L.F. (2009). Malignant astrocytomas originate from neural stem/progenitor cells in a somatic tumor suppressor mouse model. Cancer Cell 15, 45–56.

Anido, J., Saez-Borderias, A., Gonzalez-Junca, A., Rodon, L., Folch, G., Carmona, M.A., Prieto-Sanchez, R.M., Barba, I., Martinez-Saez, E., Prudkin, L., et al. (2010). TGF-beta Receptor Inhibitors Target the CD44(high)/Id1(high) Glioma-Initiating Cell Population in Human Glioblastoma. Cancer Cell 18, 655–668.

Aslakson, C.J., and Miller, F.R. (1992). Selective events in the metastatic process defined by analysis of the sequential dissemination of subpopulations of a mouse mammary tumor. Cancer Res 52, 1399–1405.

Boj, S.F., Hwang, C.I., Baker, L.A., Chio, II, Engle, D.D., Corbo, V., Jager, M., Ponz-Sarvise, M., Tiriac, H., Spector, M.S., et al. (2015). Organoid models of human and mouse ductal pancreatic cancer. Cell 160, 324–338.

Barone, M.V., Pepperkok, R., Peverali, F.A., and Philipson, L. (1994). Id proteins control growth induction in mammalian cells. Proc Natl Acad Sci USA 91, 4985–4988.

Bensellam, M., Montgomery, M.K., Luzuriaga, J., Chan, J.Y., and Laybutt, D.R. (2015). Inhibitor of differentiation proteins protect against oxidative stress by regulating the antioxidant-mitochondrial response in mouse beta cells. Diabetologia 58, 758–770.

Bhattacharya, A., and Baker, N.E. (2011). A network of broadly expressed HLH genes regulates tissue-specific cell fates. Cell 147, 881–892.

Bounpheng, M.A., Dimas, J.J., Dodds, S.G., and Christy, B.A. (1999). Degradation of Id proteins by the ubiquitin-proteasome pathway. Faseb J 13, 2257–2264.

Castanon, E., Bosch-Barrera, J., Lopez, I., Collado, V., Moreno, M., Lopez-Picazo, J.M., Arbea, L., Lozano, M.D., Calvo, A., and Gil-Bazo, I. (2013). Id1 and Id3 co-expression correlates with clinical outcome in stage III-N2 non-small cell lung cancer patients treated with definitive chemoradiotherapy. J Transl Med 11, 13.

Ciarrocchi, A., Jankovic, V., Shaked, Y., Nolan, D.J., Mittal, V., Kerbel, R.S., Nimer, S.D., and Benezra, R. (2007). Id1 restrains p21 expression to control endothelial progenitor cell formation. PLoS ONE 2, e1338.

Ciarrocchi, A., Piana, S., Valcavi, R., Gardini, G., and Casali, B. (2011). Inhibitor of DNA binding-1 induces mesenchymal features and promotes invasiveness in thyroid tumour cells. European journal of cancer (Oxford, England : 1990) 47, 934–945.

Connerney, J., Andreeva, V., Leshem, Y., Muentener, C., Mercado, M.A., and Spicer, D.B. (2006). Twist1 dimer selection regulates cranial suture patterning and fusion. Dev Dyn 235, 1345–1357.

Cook, P.J., Thomas, R., Kingsley, P.J., Shimizu, F., Montrose, D.C., Marnett, L.J., Tabar, V.S., Dannenberg, A.J., and Benezra, R. (2016). Cox-2-derived PGE2 induces Id1-dependent radiation resistance and self-renewal in experimental glioblastoma. Neuro Oncol 18, 1379–1389.

Coppe, J.P., Itahana, Y., Moore, D.H., Bennington, J.L., and Desprez, P.Y. (2004). Id-1 and Id-2 proteins as molecular markers for human prostate cancer progression. Clin Cancer Res 10, 2044–2051.

de Candia, P., Solit, D.B., Giri, D., Brogi, E., Siegel, P.M., Olshen, A.B., Muller, W.J., Rosen, N., and Benezra, R. (2003). Angiogenesis impairment in Id-deficient mice cooperates with an Hsp90 inhibitor to completely suppress HER2/neu-dependent breast tumors. Proc Natl Acad Sci U S A 100, 12337–12342.

Deed, R.W., Armitage, S., and Norton, J.D. (1996). Nuclear localization and regulation of Id protein through an E protein-mediated chaperone mechanism. J Biol Chem 271, 23603–23606.

Ding, R., Han, S., Lu, Y., Guo, C., Xie, H., Zhang, N., Song, Z., Cai, L., Liu, J., and Dou, K. (2010). Overexpressed Id-1 is associated with patient prognosis and HBx expression in hepatitis B virus-related hepatocellular carcinoma. Cancer biology & therapy 10, 299–307.

Fellmann, C., Hoffmann, T., Sridhar, V., Hopfgartner, B., Muhar, M., Roth, M., Lai, D.Y., Barbosa, I.A., Kwon, J.S., Guan, Y., et al. (2013). An optimized microRNA backbone for effective single-copy RNAi. Cell reports 5, 1704–1713.

Forootan, S.S., Wong, Y.-C., Dodson, A., Wang, X., Lin, K., Smith, P.H., Foster, C.S., and Ke, Y. (2007). Increased Id-1 expression is significantly associated with poor survival of patients with prostate cancer. Human pathology 38, 1321–1329.

Gao, D., Nolan, D.J., Mellick, A.S., Bambino, K., McDonnell, K., and Mittal, V. (2008). Endothelial progenitor cells control the angiogenic switch in mouse lung metastasis. Science 319, 195–198.

Gautschi, O., Tepper, C.G., Purnell, P.R., Izumiya, Y., Evans, C.P., Green, T.P., Desprez, P.Y., Lara, P.N., Gandara, D.R., Mack, P.C., et al. (2008). Regulation of Id1 expression by SRC: implications for targeting of the bone morphogenetic protein pathway in cancer. Cancer Res 68, 2250–2258.

Granot, Z., Henke, E., Comen, E.A., King, T.A., Norton, L., and Benezra, R. (2011). Tumor entrained neutrophils inhibit seeding in the premetastatic lung. Cancer cell 20, 300–314.

Gupta, G.P., Perk, J., Acharyya, S., de Candia, P., Mittal, V., Todorova-Manova, K., Gerald, W.L., Brogi, E., Benezra, R., and Massague, J. (2007). ID genes mediate tumor reinitiation during breast cancer lung metastasis. Proc Natl Acad Sci U S A 104, 19506–19511.

Han, S., Gou, C., Hong, L., Liu, J., ZheyiHan, Liu, C., Wang, J., Wu, K., Ding, J., and Fan, D. (2004). Expression and significances of Id1 helix-loop-helix protein overexpression in gastric cancer. Cancer letters 216, 63–71.

Hara, E., Yamaguchi, T., Nojima, H., Ide, T., Campisi, J., Okayama, H., and Oda, K. (1994). Id-related genes encoding helix-loop-helix proteins are required for G1 progression and are repressed in senescent human fibroblasts. J Biol Chem 269, 2139–2145.

Huang, Y.H., Hu, J., Chen, F., Lecomte, N., Basnet, H., David, C.J., Witkin, M.D., Allen, P.J., Leach, S.D., Hollmann, T.J., et al. (2019). ID1 mediates escape from TGF-beta tumor suppression in pancreatic cancer. Cancer Discov.

Jones, S., Zhang, X., Parsons, D.W., Lin, J.C., Leary, R.J., Angenendt, P., Mankoo, P., Carter, H., Kamiyama, H., Jimeno, A., et al. (2008). Core signaling pathways in human pancreatic cancers revealed by global genomic analyses. Science 321, 1801–1806.

Jankovic, V., Ciarrocchi, A., Boccuni, P., DeBlasio, T., Benezra, R., and Nimer, S.D. (2007). Id1 restrains myeloid commitment, maintaining the self-renewal capacity of hematopoietic stem cells. Proc Natl Acad Sci U S A 104, 1260–1265.

Kang, Y., Chen, C.R., and Massague, J. (2003). A self-enabling TGFbeta response coupled to stress signaling: Smad engages stress response factor ATF3 for Id1 repression in epithelial cells. Mol Cell 11, 915–926.

Kebebew, E., Peng, M., Treseler, P.A., Clark, O.H., Duh, Q.-Y., Ginzinger, D., and Miner, R. (2004). Id1 gene expression is up-regulated in hyperplastic and neoplastic thyroid tissue and regulates growth and differentiation in thyroid cancer cells. J Clin Endocrinol Metab 89, 6105–6111.

Kim, M.-S., Park, T.-I., Lee, Y.-M., Jo, Y.-M., and Kim, S. (2013). Expression of Id-1 and VEGF in non-small cell lung cancer. Int J Clin Exp Pathol 6, 2102–2111.

Lasorella, A., Benezra, R., and Iavarone, A. (2014). The ID proteins: master regulators of cancer stem cells and tumour aggressiveness. Nat Rev Cancer 14, 77–91.

Lasorella, A., Noseda, M., Beyna, M., and Iavarone, A. (2000). Id2 is a retinoblastoma protein target and mediates signalling by Myc oncoproteins. Nature 407, 592–598.

Le Jossic, C., Ilyin, G.P., Loyer, P., Glaise, D., Cariou, S., and Guguen-Guillouzo, C. (1994). Expression of helix-loop-helix factor Id-1 is dependent on the hepatocyte proliferation and differentiation status in rat liver and in primary culture. Cancer Res 54, 6065–6068.

Lee, K.T., Lee, Y.W., Lee, J.K., Choi, S.H., Rhee, J.C., Paik, S.S., and Kong, G. (2004). Overexpression of Id-1 is significantly associated with tumour angiogenesis in human pancreas cancers. Br J Cancer 90, 1198–1203.

Lee, S.B., Frattini, V., Bansal, M., Castano, A.M., Sherman, D., Hutchinson, K., Bruce, J.N., Califano, A., Liu, G., Cardozo, T., et al. (2016). An ID2-dependent mechanism for VHL inactivation in cancer. Nature 529, 172–177.

Li, X., Zhang, Z., Xin, D., Chua, C.W., Wong, Y.C., Leung, S.C.L., Na, Y., and Wang, X. (2007). Prognostic significance of Id-1 and its association with EGFR in renal cell cancer. Histopathology 50, 484–490.

Ling, F., Kang, B., and Sun, X.H. (2014). Id proteins: small-molecules, mighty regulators. Curr Top Dev Biol 110, 189–216.

Liu, J., Hu, Y., Hu, W., Xie, X., Ela Bella, A., and Fu, J. (2010). Expression and prognostic relevance of Id1 in stage III esophageal squamous cell carcinoma. Cancer Biomark 8, 67–72.

Liu, K.J., and Harland, R.M. (2003). Cloning and characterization of Xenopus Id4 reveals differing roles for Id genes. Dev Biol 264, 339–351.

Lyden, D., Young, A.Z., Zagzag, D., Yan, W., Gerald, W., O’Reilly, R., Bader, B.L., Hynes, R.O., Zhuang, Y., Manova, K., et al. (1999). Id1 and Id3 are required for neurogenesis, angiogenesis and vascularization of tumour xenografts [see comments]. Nature 401, 670–677.

Matsuda, Y., Yamagiwa, S., Takamura, M., Honda, Y., Ishimoto, Y., Ichida, T., and Aoyagi, Y. (2005). Overexpressed Id-1 is associated with a high risk of hepatocellular carcinoma development in patients with cirrhosis without transcriptional repression of p16. Cancer 104, 1037–1044.

Maw, M.K., Fujimoto, J., and Tamaya, T. (2008). Expression of the inhibitor of DNA-binding (ID)-1 protein as an angiogenic mediator in tumour advancement of uterine cervical cancers. British journal of cancer 99, 1557–1563.

Nam, H.S., and Benezra, R. (2009). High levels of Id1 expression define B1 type adult neural stem cells. Cell Stem Cell 5, 515–526.

Nieto, M.A., Huang, R.Y., Jackson, R.A., and Thiery, J.P. (2016). Emt: 2016. Cell 166, 21–45.

Niola, F., Zhao, X., Singh, D., Sullivan, R., Castano, A., Verrico, A., Zoppoli, P., Friedmann-Morvinski, D., Sulman, E., Barrett, L., et al. (2013). Mesenchymal high-grade glioma is maintained by the ID-RAP1 axis. J Clin Invest 123, 405–417.

Ogata, T., Wozney, J.M., Benezra, R., and Noda, M. (1993). Bone morphogenetic protein 2 transiently enhances expression of a gene, Id (inhibitor of differentiation), encoding a helix-loop-helix molecule in osteoblast-like cells. Proc Natl Acad Sci U S A 90, 9219–9222.

Pastushenko, I., Brisebarre, A., Sifrim, A., Fioramonti, M., Revenco, T., Boumahdi, S., Van Keymeulen, A.,Brown, D., Moers, V., Lemaire, S., et al. (2018). Identification of the tumour transition states occurring during EMT. Nature 556, 463–468.

Perk, J., Gil-Bazo, I., Chin, Y., de Candia, P., Chen, J.J., Zhao, Y., Chao, S., Cheong, W., Ke, Y., Al-Ahmadie, H., et al. (2006). Reassessment of id1 protein expression in human mammary, prostate, and bladder cancers using a monospecific rabbit monoclonal anti-id1 antibody. Cancer Res 66, 10870–10877.

Perry, S.S., Zhao, Y., Nie, L., Cochrane, S.W., Huang, Z., and Sun, X.H. (2007). Id1, but not Id3, directs long-term repopulating hematopoietic stem-cell maintenance. Blood 110, 2351–2360.

Pesce, S., and Benezra, R. (1993). The loop region of the helix-loop-helix protein Id1 is critical for its dominant negative activity. Mol Cell Biol 13, 7874–7880.

Ponz-Sarvise, M., Nguewa, P.A., Pajares, M.J., Agorreta, J., Lozano, M.D., Redrado, M., Pio, R., Behrens, C., Wistuba, I.I., Garcia-Franco, C.E., et al. (2011). Inhibitor of differentiation-1 as a novel prognostic factor in NSCLC patients with adenocarcinoma histology and its potential contribution to therapy resistance. Clinical cancer research : an official journal of the American Association for Cancer Research 17, 4155–4166.

Pulaski, B.A., and Ostrand-Rosenberg, S. (2001). Mouse 4T1 breast tumor model. Curr Protoc Immunol Chapter 20, Unit 20 22.

Richter, J., Schlesner, M., Hoffmann, S., Kreuz, M., Leich, E., Burkhardt, B., Rosolowski, M., Ammerpohl, O., Wagener, R., Bernhart, S.H., et al. (2012). Recurrent mutation of the ID3 gene in Burkitt lymphoma identified by integrated genome, exome and transcriptome sequencing. Nat Genet 44, 1316–1320.

Rockman, S.P., Currie, S.A., Ciavarella, M., Vincan, E., Dow, C., Thomas, R.J., and Phillips, W.A. (2001). Id2 is a target of the beta-catenin/T cell factor pathway in colon carcinoma. J Biol Chem 276, 45113–45119.

Ruzinova, M.B., Schoer, R.A., Gerald, W., Egan, J.E., Pandolfi, P.P., Rafii, S., Manova, K., Mittal, V., and Benezra, R. (2003). Effect of angiogenesis inhibition by Id loss and the contribution of bone-marrow-derived endothelial cells in spontaneous murine tumors. Cancer Cell 4, 277–289.

Schaefer, B.M., Koch, J., Wirzbach, A., and Kramer, M.D. (2001). Expression of the helix-loop-helix protein ID1 in keratinocytes is upregulated by loss of cell-matrix contact. Exp Cell Res 266, 250–259.

Schindl, M., Oberhuber, G., Obermair, A., Schoppmann, S.F., Karner, B., and Birner, P. (2001). Overexpression of Id-1 protein is a marker for unfavorable prognosis in early-stage cervical cancer. Cancer Res 61, 5703–5706.

Schindl, M., Schoppmann, S.F., Strobel, T., Heinzl, H., Leisser, C., Horvat, R., and Birner, P. (2003). Level of Id-1 protein expression correlates with poor differentiation, enhanced malignant potential, and more aggressive clinical behavior of epithelial ovarian tumors. Clinical cancer research : an official journal of the American Association for Cancer Research 9, 779–785.

Schoppmann, S.F., Schindl, M., Bayer, G., Aumayr, K., Dienes, J., Horvat, R., Rudas, M., Gnant, M., Jakesz, R., and Birner, P. (2003). Overexpression of Id-1 is associated with poor clinical outcome in node negative breast cancer. International journal of cancer 104, 677–682.

Sharma, P., Patel, D., and Chaudhary, J. (2012). Id1 and Id3 expression is associated with increasing grade of prostate cancer: Id3 preferentially regulates CDKN1B. Cancer Med 1, 187–197.

Shibue, T., and Weinberg, R.A. (2017). EMT, CSCs, and drug resistance: the mechanistic link and clinical implications. Nature reviews Clinical oncology 14, 611–629.

Stankic, M., Pavlovic, S., Chin, Y., Brogi, E., Padua, D., Norton, L., Massague, J., and Benezra, R. (2013). TGF-beta-Id1 signaling opposes Twist1 and promotes metastatic colonization via a mesenchymal-to-epithelial transition. Cell reports 5, 1228–1242.

Takahashi, M., Fukuda, K., Sugimura, T., and Wakabayashi, K. (1998). Beta-catenin is frequently mutated and demonstrates altered cellular location in azoxymethane-induced rat colon tumors. Cancer Res 58, 42–46.

Takai, N., Miyazaki, T., Fujisawa, K., Nasu, K., and Miyakawa, I. (2001). Id1 expression is associated with histological grade and invasive behavior in endometrial carcinoma. Cancer letters 165, 185–193.

Takai, N., Ueda, T., Nishida, M., Nasu, K., and Miyakawa, I. (2004). The relationship between oncogene expression and clinical outcome in endometrial carcinoma. Curr Cancer Drug Targets 4, 511–520.

Tang, R., Hirsch, P., Fava, F., Lapusan, S., Marzac, C., Teyssandier, I., Pardo, J., Marie, J.-P., and Legrand, O. (2009). High Id1 expression is associated with poor prognosis in 237 patients with acute myeloid leukemia. Blood 114, 2993–3000.

Tournay, O., and Benezra, R. (1996). Transcription of the dominant-negative helix-loop-helix protein Id1 is regulated by a protein complex containing the immediate-early response gene Egr-1. Mol Cell Biol 16, 2418–2430.

Tsang, C.M., Cheung, K.C., Cheung, Y.C., Man, K., Lui, V.W., Tsao, S.W., and Feng, Y. (2015). Berberine suppresses Id-1 expression and inhibits the growth and development of lung metastases in hepatocellular carcinoma. Biochim Biophys Acta 1852, 541–551.

Vandeputte, D.A., Troost, D., Leenstra, S., Ijlst-Keizers, H., Ramkema, M., Bosch, D.A., Baas, F., Das, N.K., and Aronica, E. (2002). Expression and distribution of id helix-loop-helix proteins in human astrocytic tumors. Glia 38, 329–338.

Wang, L.H., and Baker, N.E. (2015). E Proteins and ID Proteins: Helix-Loop-Helix Partners in Development and Disease. Dev Cell 35, 269–280.

Wazir, U., Jiang, W.G., Sharma, A.K., Newbold, R.F., and Mokbel, K. (2013). The mRNA expression of inhibitors of DNA binding-1 and -2 is associated with advanced tumour stage and adverse clinical outcome in human breast cancer. Anticancer Res 33, 2179–2183.

Welsch, M.E., Kaplan, A., Chambers, J.M., Stokes, M.E., Bos, P.H., Zask, A., Zhang, Y., Sanchez-Martin, M., Badgley, M.A., Huang, C.S., et al. (2017). Multivalent Small-Molecule Pan-RAS Inhibitors. Cell 168, 878–889 e829.

Wen, Y.H., Ho, A., Patil, S., Akram, M., Catalano, J., Eaton, A., Norton, L., Benezra, R., and Brogi, E. (2012). Id4 protein is highly expressed in triple-negative breast carcinomas: possible implications for BRCA1 downregulation. Breast Cancer Res Treat 135, 93–102.

Wojnarowicz, P.M., Lima, E.S.R., Ohnaka, M., Lee, S.B., Chin, Y., Kulukian, A., Chang, S.H., Desai, B., Garcia Escolano, M., Shah, R., et al. (2019). A small-molecule pan-Id antagonist inhibits pathologic ocular neovascularization. Cell Reports 29, 62–75 e67.

Yang, H.-Y., Liu, H.-L., Liu, G.-Y., Zhu, H., Meng, Q.-W., Qu, L.-D., Liu, L.-X., and Jiang, H.-C. (2011). Expression and prognostic values of Id-1 and Id-3 in gastric adenocarcinoma. The Journal of surgical research 167, 258–266.

Ying, Q.L., Nichols, J., Chambers, I., and Smith, A. (2003). BMP induction of Id proteins suppresses differentiation and sustains embryonic stem cell self-renewal in collaboration with STAT3. Cell 115, 281–292.

Zhang, N., Subbaramaiah, K., Yantiss, R.K., Zhou, X.K., Chin, Y., Benezra, R., and Dannenberg, A.J. (2015). Id1 Deficiency Protects against Tumor Formation in ApcMin/+ Mice but Not in a Mouse Model of Colitis-Associated Colon Cancer. Cancer prevention research 8, 303–311.

Zhang, N., Yantiss, R.K., Nam, H.S., Chin, Y., Zhou, X.K., Scherl, E.J., Bosworth, B.P., Subbaramaiah, K., Dannenberg, A.J., and Benezra, R. (2014). ID1 is a functional marker for intestinal stem and progenitor cells required for normal response to injury. Stem cell reports 3, 716–724.

Zhao, Z.-R., Zhang, Z.-Y., Zhang, H., Jiang, L., Wang, M.-W., and Sun, X.-F. (2008). Overexpression of Id-1 protein is a marker in colorectal cancer progression. Oncology reports 19, 419–424.

Zimel, M.N., Horowitz, C.B., Rajasekhar, V.K., Christ, A.B., Wei, X., Wu, J., Wojnarowicz, P.M., Wang, D., Goldring, S.R., Purdue, P.E., et al. (2017). HPMA-Copolymer Nanocarrier Targets Tumor-Associated Macrophages in Primary and Metastatic Breast Cancer. Mol Cancer Ther 16, 2701–2710.

